# Discovery of interpretable patterning rules by integrating mechanistic modeling and deep learning

**DOI:** 10.1101/2024.09.02.610872

**Authors:** Jia Lu, Nan Luo, Sizhe Liu, Kinshuk Sahu, Rohan Maddamsetti, Yasa Baig, Lingchong You

**Author notes:** Corresponding author. Department of Biomedical Engineering, Duke University, CIEMAS 2355, 101 Science Drive, Box 3382, Durham, NC 27708, USA.

## Abstract

Predictive programming of self-organized pattern formation using living cells is challenging in major part due to the difficulty in navigating through the high-dimensional design space effectively. The emergence and characteristics of patterns are highly sensitive to both system and environmental parameters. Often, the optimal conditions able to generate patterns represent a small fraction of the possible design space. Furthermore, the experimental generation and quantification of patterns is typically labor intensive and low throughput, making it impractical to optimize pattern formation solely based on trials and errors. To this end, simulations using a well-formulated mechanistic model can facilitate the identification of optimal experimental conditions for pattern formation. However, even a moderately complex system can make these simulations computationally prohibitive when applied to a large parameter space. In this study, we demonstrate how integrating mechanistic modeling with machine learning can significantly accelerate the exploration of design space for patterning circuits and aid in deriving human-interpretable design rules. We apply this strategy to program self-organized ring patterns in *Pseudomonas aeruginosa* using a synthetic gene circuit. Our approach involved training a neural network with simulated data to predict pattern formation 10 million times faster than the mechanistic model. This neural network was then used to predict pattern formation across a vast array of parameter combinations, far exceeding the size of the training dataset and what was computationally feasible using the mechanistic model alone. By doing so, we identified many parameter combinations able to generate desirable patterns, which still represent an extremely small fraction of explored parametric space. We next used the mechanistic model to validate top candidates and identify coarse-grained rules for patterning. We experimentally demonstrated the generation and control of patterning guided by the learned rules. Our work highlights the effectiveness in integrating mechanistic modeling and machine learning for rational engineering of complex dynamics in living cells.

## Introduction

Pattern formation is essential to the arrangement and organization of biomolecules, cells, and tissues that define the structure and function of diverse organisms or systems, at a wide range of length scales (from sub-microns to kilometers). Examples include the formation of the division ring in a bacterial cell^1^, the development of embryonic cells^2^, the growth of bacterial colonies^3^, as well as the organization of macroscopic ecosystems^4^.

A major challenge facing the study of pattern formation using natural biological systems (e.g. development of animal body plan) is the incredible complexity of the interactions underlying such processes. The emergence of modern synthetic biology offers an alternative with an appealing advantage. By removing confounding factors, predicting and controlling pattern formation in a reduced system can offer insights into the essential requirement of generating certain self-organized patterns. Moreover, the ability to do so provides a strategy to engineer living structures for practical applications. Indeed, there have been substantial efforts in rational programming of spatial patterns since the inception of this field of research^5–16^.

Even when we have complete control on the choice of cellular components, our ability to generate patterns in a *predictive* manner is extremely limited, in comparison to our ability to program other types of dynamics (e.g., logic gates)^11,17^. Of the numerous gene circuits built to date, ~10 circuits have been demonstrated to generate self-organized pattern formation at population level^5–9,13–15,18^. The difficulty in predictive programming of self-organized patterns is due to several factors. First, it is possible that robust pattern formation observed in nature may depend on many of the “confounding” factors removed in the design of gene circuits. Even though a simplified, engineered system can generate patterns, the conditions to do so become much more constraining, making it difficult to achieve experimentally. For instance, while the Turing mechanism that can in theory lead to diverse patterns in a deterministic manner, it has not been conclusively demonstrated in synthetic biology, in part because the target parameter space for pattern formation is narrow and the resulting patterns are highly sensitive to perturbations. As a result, experimentally achieving the precise control of each part is difficult^19,20^.

Second, predicting pattern formation by simulation poses a unique challenge, in comparison to other types of dynamics. Prediction of pattern formation requires computation of dynamics in both time and space (using partial differential equations or agent-based models), which makes it computationally demanding and much less intuitive. Even if a circuit in theory can generate certain patterning dynamics, it is challenging to quickly identify the biologically plausible parameter ranges to realize this ability.

To this end, we present a generalizable approach, which integrates mechanistic modeling and machine learning, to quickly navigate through the vast parameter space, enabling us to deduce coarse-grained rules to guide experimental design. A key element for acceleration was the incorporation of a generative machine learning (ML) model. Generative models can make reliable predictions by learning from training data and predicting from the underlying distributions. Here, we trained a ML model on simulation data from a partial differential equation (PDE) mechanistic model. The trained ML model can generate predictions much faster but lacks the capacity for direct interpretation of the predictions. As such, we validated the ML model performance with top parameter combinations against numerical solutions. This approach maximally leveraged the prior biological knowledge while exploiting the computational acceleration offered by ML.

As a demonstration, we applied this approach to guide the porting of a synthetic patterning circuit, previously demonstrated in *E. coli* ^15^, into a new host -- *Pseudomonas aeruginosa* (PA14). The new host, *P. aeruginosa*, differs from *E. coli* in several traits, including growth rate, biosurfactant production, and cell motility^21,22^. When grown on a semi-solid surface and starved, it can form branching patterns, which are the collective results of growth, cell motility, and biosurfactant induced surface tension gradient. It also has multiple hierarchical quorum sensing (QS) systems, adding additional possibilities for interactions and crosstalk. The difference in the host physiology poses two technical challenges in porting the circuit and optimizing its dynamics. First, can the ring-forming capability be maintained in the new host, given a greater degree of interference from the host QS system? Second, if so, does the presence of additional chassis capability enable robust generation of more complex patterns than previously demonstrated?

Using our integrated approach, we systematically examined the vast parameter space for the circuit dynamics in the new host and derived and validated coarse-grained rules for generating complex patterns. The rules highlight the key requirement on circuit dynamics, as well as the contribution of host physiology and environmental conditions. These criteria pointed us to the experimental conditions for generating up to five rings, greatly exceeding the two rings in *E. coli*. Furthermore, we used seeding array with bioprinting tool, to show that the reaction-diffusion mechanism can be superimposed on top of pre-defined patterns or symmetry breaking dynamics. It shows that compositional mechanisms is a promising engineering approach in creating novel, complex spatial organization.

## Results

### Porting a patterning circuit into *P. aeruginosa*

The new patterning system consists of two parts – a reaction-diffusion gene circuit (Figure 1A), and a PA14 host (Figure 1B). In the circuit, T7 RNA polymerase (T7RNAP) serves as the activator by activating its own expression and that of T7 lysozyme. T7 lysozyme is the inhibitor which binds with T7RNAP forming a complex and inhibits T7RNAP expression through repressing the upstream T7 promoter^23^. 3OC6HSL is a quorum sensing (QS) small molecule that freely diffuses in agar and in colony, connecting the activator and inhibitor nodes of the circuit. Figure 1C illustrates the experimental procedure for porting the circuit to PA14. To allow circuit function and maintenance in PA14, we experimented a series of optimization on plasmid design, including the choice of shuttle vector, replication origin, antibiotic resistance marker, and fluorescent reporters. Due to PA14’s natural resistance to a wide range of antibiotics, strong autofluorescence, limited available replication origins, and circuit’s high metabolic burden, we focused on a construct, in which a host-specific replication origin pRO1600 fragment was introduced in addition to p15A origin, gentamycin resistance gene was used for selection, and the entire gene circuit was encoded on a single plasmid (Supplementary figure 1)^24^. The plasmid copy number in PA14 appeared to be low (∼3 copies/cell).

**Figure 1.**
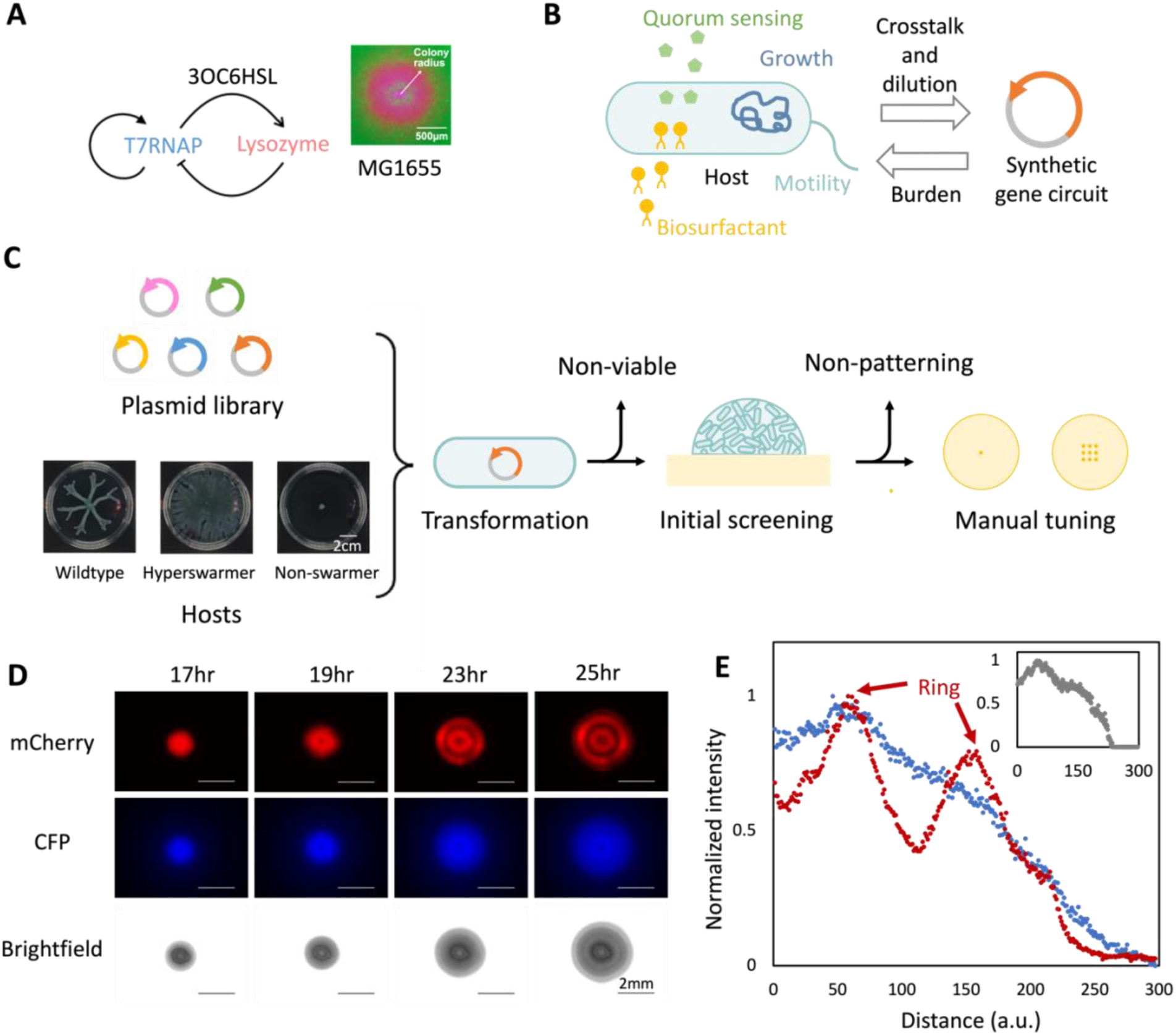
Construction of the PA14 patterning system. The system is a core-ring patterning circuit (previously developed in *E. coli*) ported into a PA14 host. a. The patterning gene circuit. T7 RNAP is the activator and T7 lysozyme is the inhibitor. They are co-expressed with CFP and mCherry, respectively. The inset shows a core-ring pattern (mCherry channel) formed in *E. coli* MG1655 (reproduced from Cao *et al.*^29^ with permission). b. PA14 host. PA14 has unique characteristics in terms of growth, motility, and endogenous quorum sensing systems, which will influence circuit dynamics. c. Experimental procedure for constructing the new patterning system. The circuit was modified into different designs to adapt to PA14 hosts, and they were transformed into PA14 hosts with different colony morphology. After electroporation and selecting on antibiotic selection plates, the viable systems were used for patterning experiments, where cells were manual pipetted onto agar plate with chemical inducer IPTG. The ones that could express fluorescent signals and form patterns were then bio-printed using a liquid handling system, with different volumes or in array, to test their patterning capacities under well-controlled conditions (see Methods). d. Time course of pattern development in PA14-*lasR*. 0.1 μl of OD = 0.2 cell culture was inoculated at the center of an agar plate and incubated at 37°C. The agar plate contained 0.5% agar (w/v), 10 g/L casamino acid, 1 mM IPTG, and 10 μg/ml gentamycin. e. Quantification of colony patterns by cross profiling. As the colonies are approximately circular, cross profiles of the fluorescent and brightfield channels were measured from colony center to edge. The intensities were normalized according to their respective maximum intensities. This profile represents the colony at 25hr from Fig 1D. Two mCherry peaks indicates a two-ring pattern. Red: mCherry, blue: CFP, grey: cell density.

When selecting host strains, colony phenotype was considered as they are indicators of host physiology, such as motility, growth rate, flagella movement, and QS systems^25^. We transformed the plasmid into PA14 strains with three distinct swarming phenotypes by electroporation^26^. The wildtype PA14 and a *fleN* mutant (hyperswarmer)^27^ could not maintain the circuit; however, a *lasR* mutant (PA14-*lasR*)^25^ could stably maintain the circuit. This difference was likely due to the difference in the strength of QS systems in these different strains. The WT and the *fleN* mutant have intact *las* QS system, and its 3OC12HSL binds with LasR or LuxR, enhancing plux promoter activation^28^. This upregulates the circuit’s mCherry expressions and makes it burdensome, even without further induction. In contrast, the PA14-*lasR* exhibited reduced the *las-lux* promoter crosstalk^28^. Unless noted otherwise, all our subsequent experiments were carried out using the *lasR* mutant.

When inoculating the cell culture on agar surface, the PA14-*lasR* system exhibited similar behavior as the *E. coli* counterpart -- a core-ring pattern formed in mCherry channel over a day (Figure 1D). The pattern was evident from the cross profiles of fluorescent and brightfield channels as well (Figure 1E). The pattern formation also responded to exogenous IPTG and 3OC6HSL both on agar and in liquid culture (Supplementary figure 2-3). Adding IPTG shifted the circuit dynamic towards positive feedback, promoting ring formation; adding 3OC6HSL enhanced the negative feedback weakening the patterns; both findings agreeing with previous observations in *E. coli* ^29^, demonstrating the gene circuit’s primary function is preserved in the new host.

### Mechanistical model capturing the pattern formation dynamics

Upon demonstrating the qualitative portability of the circuit dynamics in the new host, we wonder if it could be tuned to generate more sophisticated dynamics. In our previous studies, we observed at most 2 rings in *E. coli* MC4100Z1 and MG1655 colonies^15,29^, though computation suggests it has the potential to generate more rings^30^.

Tuning the circuit dynamics manually by trial and error would be impractical, especially with the introduction of additional host factors. To this end, we developed a pipeline integrating mechanistic model, generative ML, and quantitative experiments to quickly explore the system dynamics (Figure 2A). The first step was to develop a reliable PDE mechanistic model to describe the patterning dynamics. In this model, gene expression, cell growth and movement, nutrient diffusion and consumption, QS molecule diffusion and crosstalk were incorporated (see Supplementary information). The model permits free lateral movement of cells and small molecules, reflecting colony growth on an agar surface. Compared to our previous work with *E. coli* which produced patterns on the length scale of < 0.5mm, the patterns formed by the new system range from 1 – 5 mm, whereby the diffusion of small molecules becomes potentially important.

**Figure 2.**
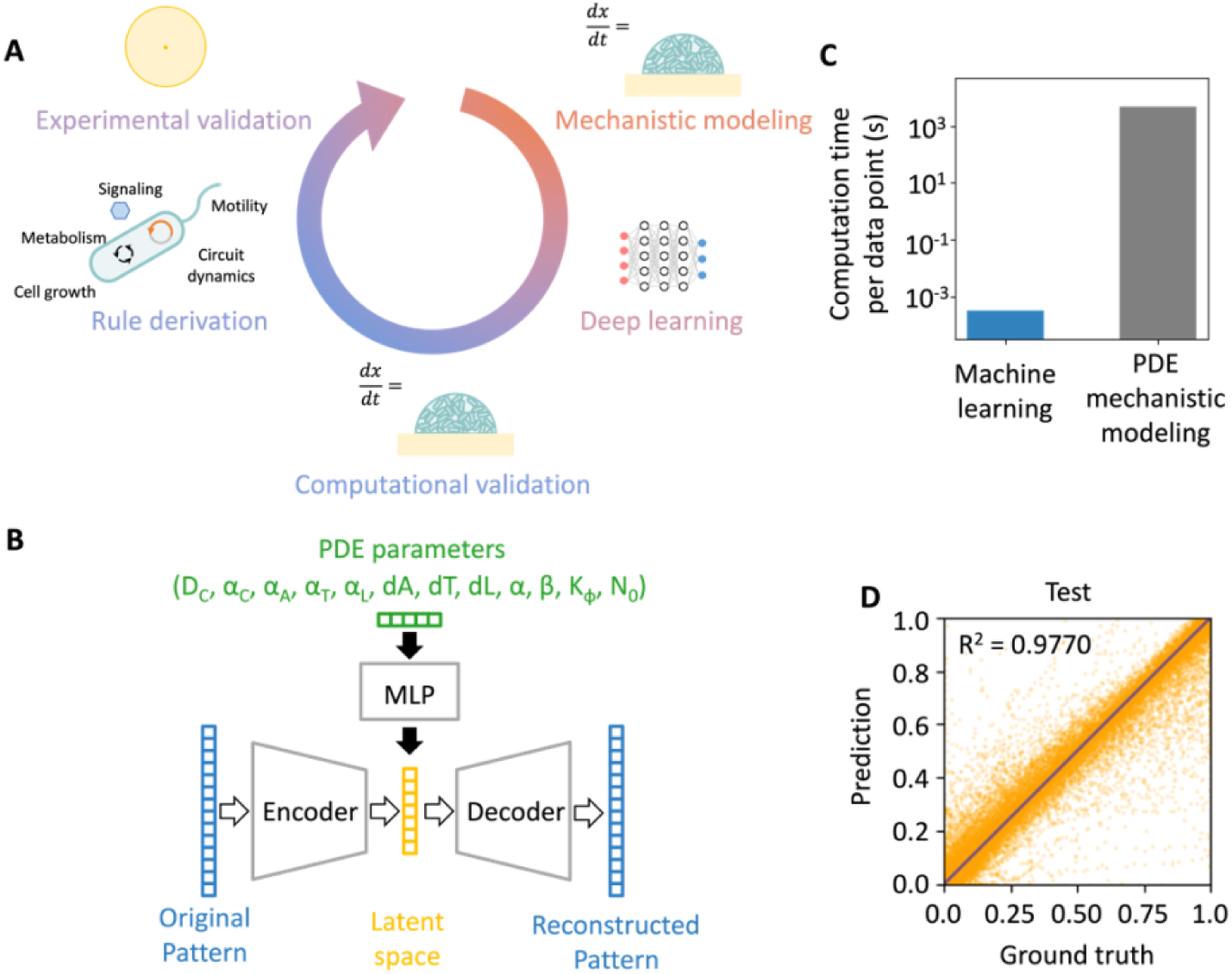
Integrating mechanistic modeling and machine learning for fast and reliable exploration of the vast parametric space of potential circuit dynamics. A. Rule derivation workflow overview. 1) Constructing a PDE mechanistic model and numerically solving it to generate a training dataset. 2) Training a machine learning (ML) surrogate model using the simulation dataset. 3) Predicting patterns from new, randomly sampled parameter sets, and validating the ML predictions against PDE simulations. 4) Constructing a dataset using the validated data; based on it, identifying key parameters for patterning and interpreting into coarse-grained rules. 5) Experimentally validating the rules by quantitatively perturbing the related experimental conditions. B. Machine learning model architecture. The model is made of a multilayer perceptron (MLP) and a variational encoder (VAE)’s decoder part. We first trained a VAE using simulation data (empty arrow). It learns to compress the 1D mCherry profiles into low-dimensional latent vectors through an encoder, then decompress them back into the original profiles through a decoder. The VAE’s training uses a reconstruction loss composed of two parts: one part that compares the reconstructed profiles to the original ones using mean squared error, and another that minimizes the Kullback-Leibler (KL) divergence between the learned latent distributions and a standard normal distribution. Next, we trained a MLP connected to the trained VAE decoder part (solid arrow). Such that it uses the 12 PDE parameters to predict the corresponding 1D mCherry profiles. The same reconstruction loss was used to update the weights of the MLP. For inference, the trained MLP and trained decoder were connected. C. Computation speed of PDE numerical simulation and ML surrogate model prediction. D. ML surrogate performance on test set. 2, 000 random samples were drawn from the test set for plotting, all samples from the test set were used for R^2^ calculation.

This model contains 31 parameters in total. We picked 12 to focus on based on two criteria (Supplementary table 1). First, they are likely to play an important role in pattern formation based on previous work and our pilot simulations. For instance, previous studies^29,30^ show the strength of the positive and negative feedback are critical for reaction diffusion systems outputs, thus parameters such as T7RNAP’s synthesis and degradation rates were chosen. Host context was known to impact gene expression^31–33^, thus cell growth rate and motility were also chosen. Second, the parameters should be experimentally tunable. For instance, the degradation rate of AHLs is tunable through pH, and the initial nutrient concentration is controllable through formulation of the growth media. Our goal is to use computation to deduce coarse-grained rules that will guide us to identify promising experimental conditions favoring the generation of multiple rings.

### Massive acceleration in pattern predictions by using machine learning

Depending on parameter choices and numerical configurations, each PDE simulation takes ∼15 min on average. With the speed, it is computationally prohibitive to extensively explore the parametric space defined by the 12 parameters. To address this challenge, we developed a ML model to emulate the PDE model in predicting final patterns, but at a much faster speed.

We carried out numerical simulations to generate a training dataset of 30,000 samples, each consisting of a randomly sampled parameter combination and the corresponding patterning profile. A multilayer perceptron - variational autoencoder (MLP-VAE) architecture was used due to its outstanding performance in capturing nonlinear dynamics and its simple and straightforward training process (Figure 2B). Specifically, a VAE was first trained to reconstruct the 1D pattern profiles, with high accuracy (R^2^ = 0.9955, Supplementary fig 4). Then the decoder part was frozen, and an MLP was placed in front of the decoder. In this way, the MLP learns to predict the VAE latent vector from the 12 PDE parameters, and the decoder converts the latent vectors into 1D patterns (See Methods).

With the ML surrogate, predicting patterns from unseen random parameters sets achieved a 10^7^ fold acceleration of over PDE simulation on the same CPU hardware (Figure 2C). The model performance was high – on the test set, the R^2^ was 0.9770, the accuracy on predicting pattern type was 95.93 % (Figure 2D, Supplementary figure 5).

To deduce rules that favor the generation of multiple rings, we wanted to generate sufficient samples with the ability to generate two or more rings. To this end, we sampled 16 million parameter combinations (∼533 times larger than the training set) using the ML surrogate model within seconds. From these, we chose 15,000 parameter combinations, distributed evenly with 5,000 samples each from 1, 2, and 3 ring classes, to validate against PDE simulations. The overall R^2^ was 0.6495. We consider a sample as validated if the ring number is correct or the R^2^ > 0.8. Following this criterion, most samples (56.06%) were validated as reliable (Supplementary figure 6).

### Deriving patterning rules to guide experimental optimization

The ML surrogate is critical for the discovery of coarse-grained patterning rules and parameter combinations, because the patterning parameter space is very narrow and shrinks rapidly for more rings (Supplementary figure 7). Only with the massive acceleration, we could extensively screen the parameter space.

From the validated dataset, we randomly chose a dataset of 4, 000 samples, with 2, 000 forming 1 ring, 1, 000 forming 2 rings, and 1, 000 forming 3 rings (Figure 3A-B, Supplementary figure 6) and analyzed their distributions. 4 out of 12 parameters were found critical for pattern emergence (Figure 3C). They represent different biological factors, including high motility (*D_C_*), small growth rate (α_*C*_), and strong positive feedback (α_*T*_ and *d*_*T*_). They represent not only circuit dynamics, but also host physiology and environmental conditions. Similarly, pairwise correlations between parameters were also found (Figure 3D). There is synergy between positive feedback strength and the growth rate and motility – when increasing motility or growth rate, stronger positive feedback is required for generating multiple rings. Besides, higher motility needs higher growth rate to compensate.

**Figure 3.**
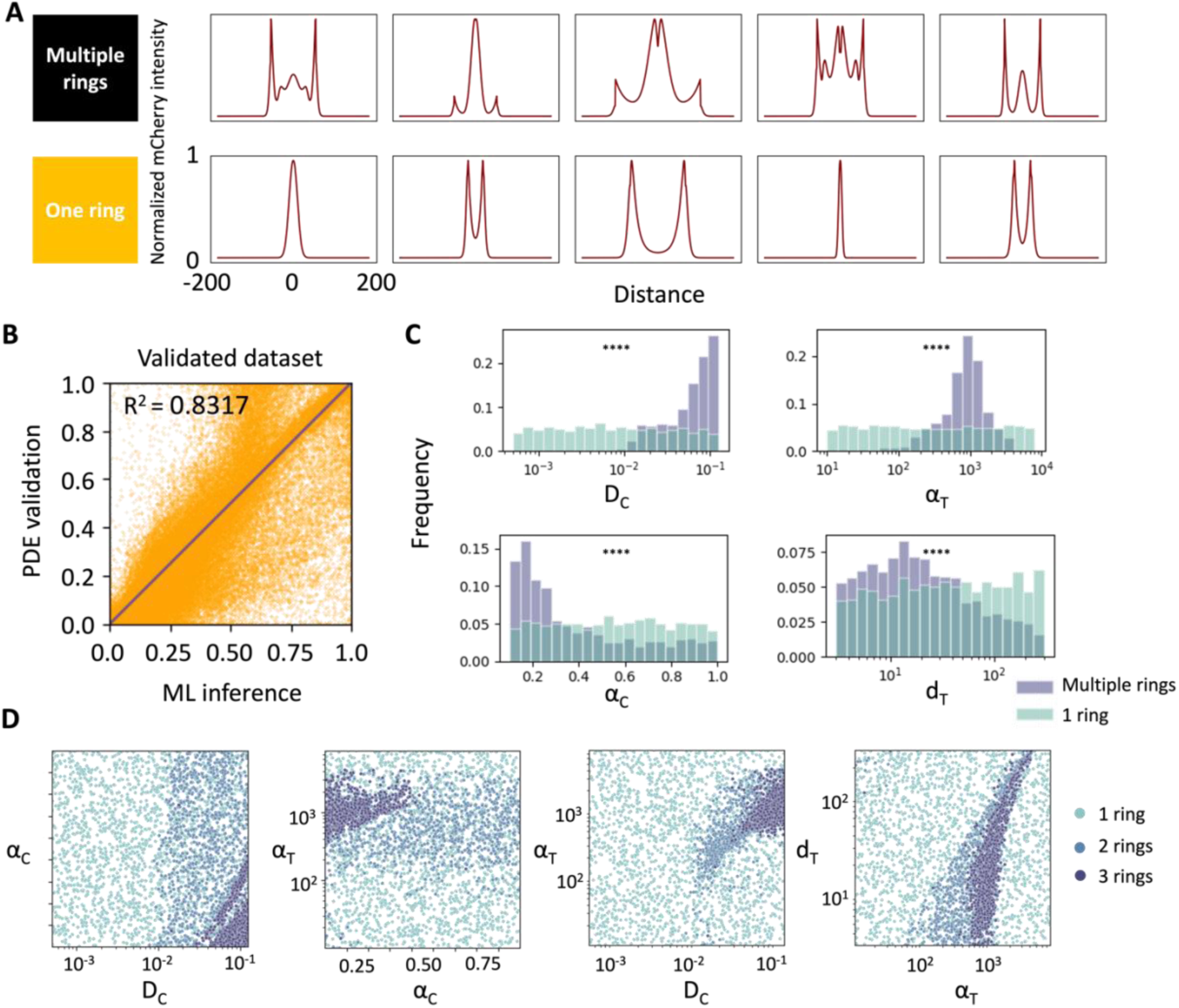
Rule derivation from ML-aided parameter screening. The dataset with 4000 validated data points was used for this analysis. A. Examples of one ring and multiple ring patterns. The ring number is identified based on the peak numbers in the full 1D mCherry cross profiles using a customized algorithm (See Methods). B. R^2^ on the final validated dataset. The dataset is made of 2, 000 1 ring patterns, 1, 000 2 ring patterns and 1, 000 3 ring patterns. The R^2^ is tested by comparing ML predictions against the PDE solutions. 2, 000 random samples were used for plotting, all 4, 000 samples were used for R^2^ calculation. The validated dataset was used for downstream analysis in C and D. Key individual parameters for pattern formation. The Mann-Whitney U test was applied to determine if the multiple ring parameter distributions were distinct from the entire dataset. **** indicates p-value < 0.0001. The other parameters were not critical for generating multiple rings (Supplementary Figure 8). C. Key pairwise correlations between the parameters for patterning.

Importantly, these factors can be controlled in experiments to validate the coarse-grained computation-derived rules (Figure 4). Agar concentration determines growth substrate stiffness and modulates cell motility. A lower agar density leads to higher motility, which is more likely to generate more rings (Figure 4A). Higher IPTG concentration enhances positive feedback and leads to more rings and deeper groove between mCherry peaks (Figure 4B). Nutrient impacts the growth rate in a biphasic manner, large or small concentration gives lower growth rate, intermediate concentration gives higher growth rate (Figure 4C). Consistent with the derived rules, both 4 g/L and 16 g/L casamino acid concentration result in deep grooves between the mCherry peaks (Figure 4D). For the pairwise rules, the combination of higher IPTG concentration and lower agar density, and the combination of higher IPTG concentration and small growth rate, both led to more and clearly separated rings (Supplementary figures 9 &10). Note that we only focused on the coarse-grained rules, because the system was intrinsically sensitive; a small noise in the parameters could result in the disappearance of patterns. These rules suggest conditions that are conducive to the emergence of complex patterns, rather than guaranteeing their formation.

**Figure 4.**
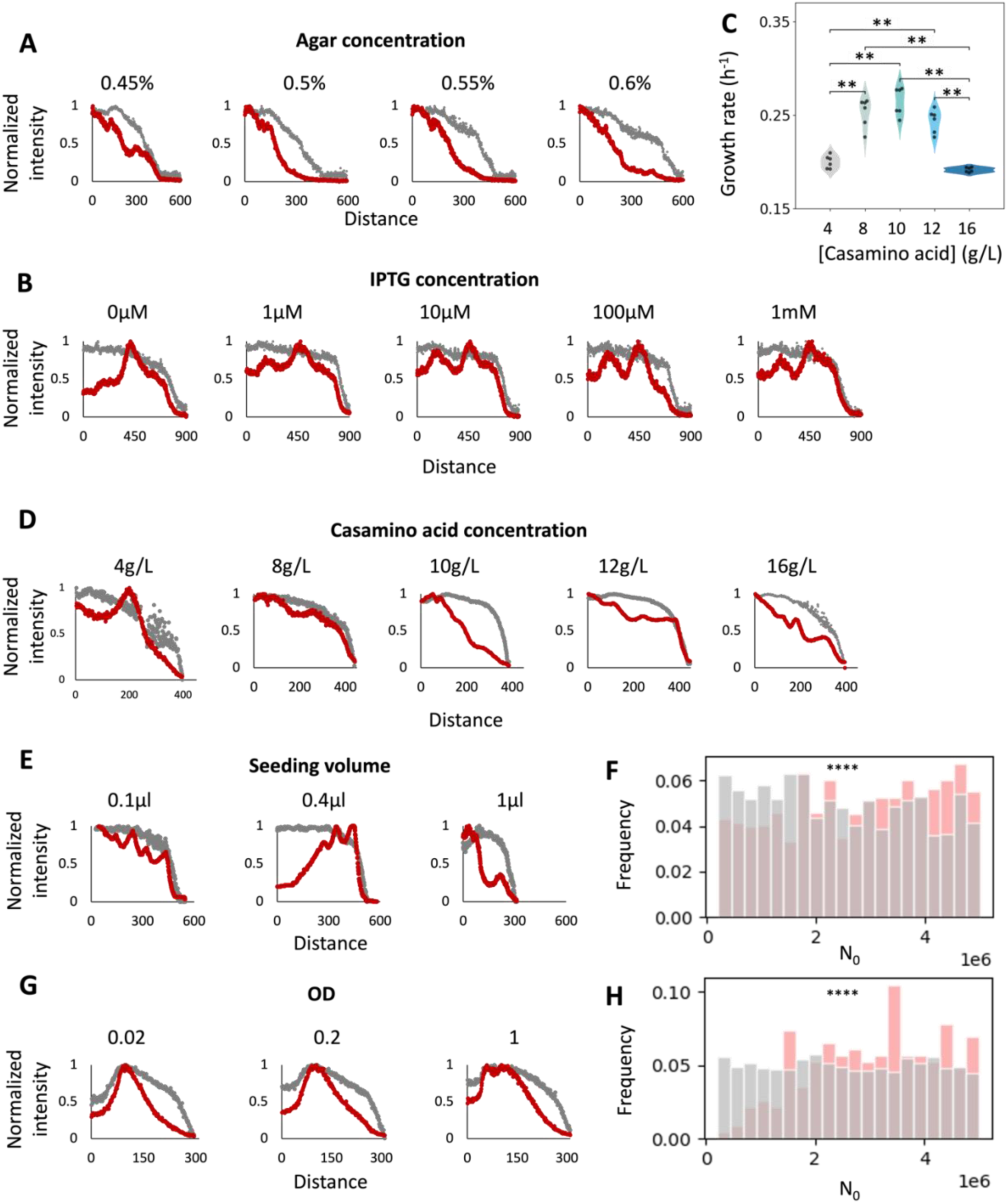
Experimental validation of computationally derived coarse-grained rules. The colony cross profiles were used to quantitatively illustrate the experimental patterns. Red and grey curves are mCherry and brightfield intensities, respectively. All conditions were kept at their default settings except for the condition being varied. A. Effects of motility. A lower agar density leads to higher cell motility, favoring ring formation B. Effects of positive feedback strength. A higher IPTG concentration leads to stronger circuit induction and stronger positive feedback, favoring ring formation. C. Nutrient concentration affects growth rate in liquid culture in a non-monotonic manner. The Mann-Whitney U test is carried out to compare if the growth rates are significantly different. ** indicates 0.01 > p-value > 0.001; if not labeled, p-value > 0.05. D. Effects of nutrient concentration. Low or high nutrient concentrations led to slower growth rates (Figure 4C), favoring ring formation. E. A smaller seeding volume led to more rings. 0.1, 0.4, and 1 μl OD = 0.2 cell culture was inoculated on agar surface and incubated for 25 hours before imaging. F. A larger nutrient concentration (*N*_0_) is important to the seeding volume dependent patterning (at fixed seeding density). The seeding density was fixed to 0.2 and the 12 PDE parameters were randomly sampled to create 3,000 different parameter combinations. For each parameter combination, 5 seeding volumes (0.1, 0.2, 0.3, 0.4, and 1 μl) were tested, and the trend was analyzed. Pink indicates parameters generating more rings with larger seeding volume, grey indicates the other cases (Supplementary figure 11A). G. A higher cell density led to more rings. 0.1 μl of OD = 0.02, 0.2, or 1 cell culture was seeded on agar surface, and incubated for 25 hours before imaging. H. Larger nutrient concentration (*N*_0_) is important to the cell density dependent patterning (at fixed seeding volume). Seeding volume was fixed to 0.1 μl and the 12 PDE parameters were randomly sampled to create 3,000 different parameter combinations. For each parameter combination, 3 seeding densities (0.02, 0.2, and 1) were tested, and the trend was analyzed. Pink indicates parameters generating more rings with higher seeding density, grey indicates the other cases (Supplementary figure 11B).

The system also exhibits seeding volume dependent patterning capabilities (Figure 4E). We used a liquid dispenser system (MANTIS, Formulatrix) to print the cell culture (OD = 0.2) onto agar surface. The volume precision was refined to 0.1 μl, and the spatial precision was achieved up to 2.25 mm. When seeding 0.1 μl cell culture, we observed four rings and a core; seeding 0. 4 μl resulted in 3 rings, and 1 μl produced 2 rings. Smaller seeding volume tended to give more rings.

The seeding volume can affect pattern formation through two parameters – the initial cell number or the starting domain size. Since growing from a few cells only led to two rings in *E.coli* ^15,29^, it is possible that the more cells led to faster colony development. By using the same method of screening (Figure 2), we investigated the contributing factor to this observation. In the mechanistic model, seeding volume was incorporated as a new parameter. The inoculum diameter scaled according to experimental measurement. When fixing seeding cell density and varying seeding volume, a small fraction of the parameter combinations exhibited the observed trend, while the rest showed the opposite trend or invariant to seeding volume (Supplementary figure 11A). Large nutrient concentration (*N*_0_) was found to be critical for this observation (Figure 4F), in addition to the other rules. On the other hand, the cell density could also play a role to the trend. Indeed, when fixing the seeding volume to 0.1 μl, higher OD led to more evident rings experimentally (Figure 4G). From parameter screening, larger nutrient concentration (*N*_0_) was also found to be critical for this observation (Figure 4H, Supplementary figure 11B).

To summarize, strong positive feedback, slow growth, large motility, large nutrient concentration, small seeding volume, and high cell density favor multiple ring generation, and there is also synergy between some of these conditions.

### Composite patterns by combining different patterning modes

The coarse-grained rules suggested experimental conditions to find extended, complex patterns with higher probability. Indeed, we were able to reproducibly discover different types of ring patterns in experiments with the favored conditions (Figure 5A).

**Figure 5.**
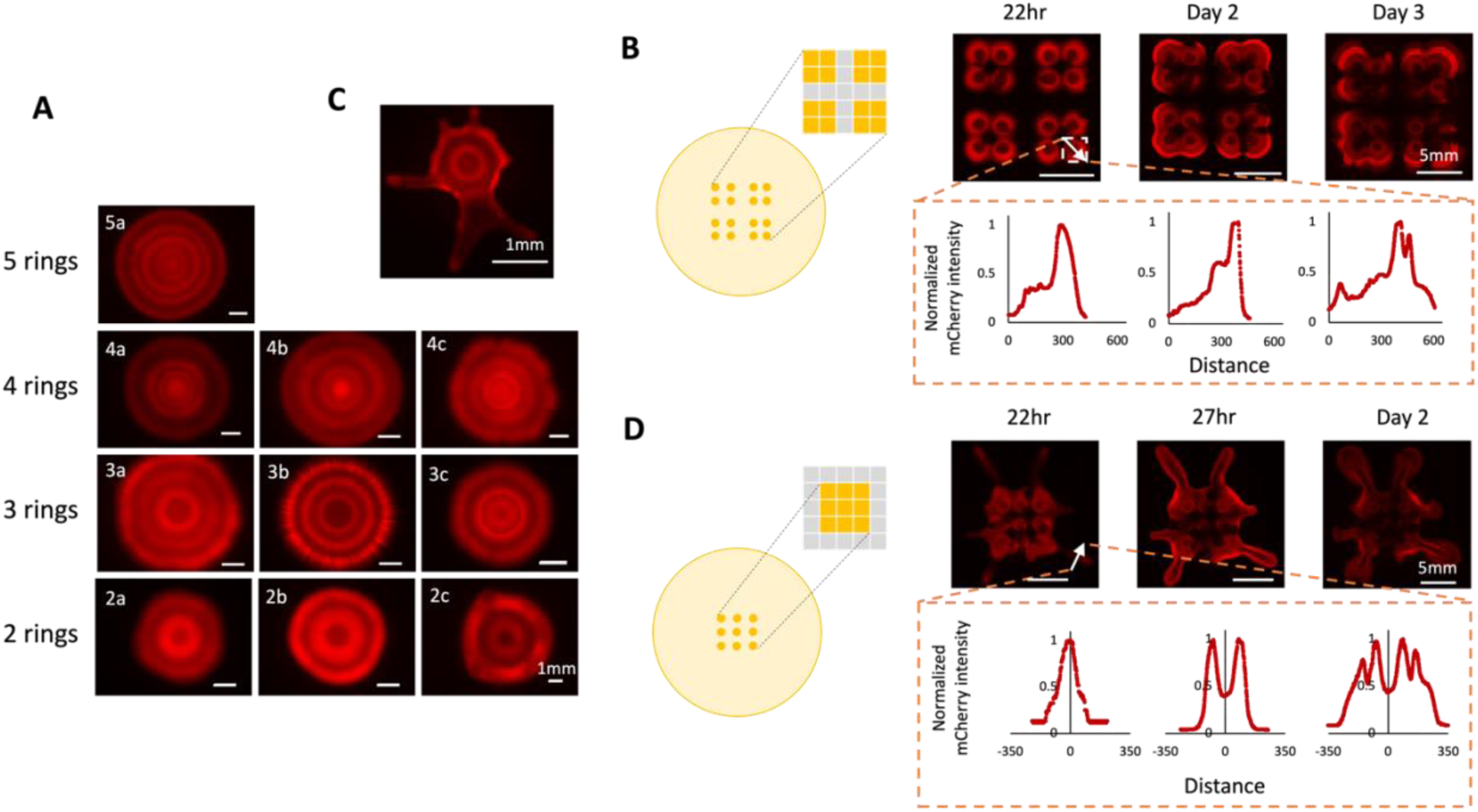
Extended pattern formation with the computational derived rules. A. Generation of multiple ring patterns in single colonies. 0.1 μl OD = 0.2 cell culture was inoculated on agar surface. The growth conditions are: 2a: 0.5% agar, 10g/L casamino acid (CA), 1mM IPTG; 2b: 0.5% agar, 8g/L CA, 1mM IPTG; 2c: 0.5% agar, 10g/L CA, 0.1mM IPTG; 3a: 0.5% agar, 10g/L CA, 1mM IPTG; 3b: 0.6% agar, 10g/L CA, 1mM IPTG; 3c: 0.55% agar, 10g/L CA, 1mM IPTG; 4a: 0.5% agar, 16/L CA, 1mM IPTG; 4b: 0.5% agar, 16g/L CA, 0mM IPTG; 4c: 0.5% agar, 16g/L CA, 1mM IPTG; 5a: 0.5% agar, 8g/L CA, 1mM IPTG. B. Composite patterns driven by self-organization and pre-patterning. Cell culture was bio-printed into an array on agar surface. The array organization is shown in the colored table, yellow indicates seeded with 0.1 μl OD = 0.2 cell culture, grey indicates no seeding. The two immediately neighboring blocks have a center-to-center distance of 2.25 mm, each inoculum has a diameter of 0.88 mm. A colony on the outer edge of the array was selected (white box) for pattern quantification. The cross profile was measured along the white arrow. C. Composite branching – ring pattern formed in a single colony. D. Composite branching – ring patterns with seeding array. Cell culture was bio-printed into a 3×3 array on agar surface. The array organization is shown in the colored table, yellow indicates seeded with 0.1 μl OD = 0.2 cell culture, grey indicates no seeding. The center-to-center distance between two neighboring inoculum is 2.25 mm, each inoculum has a diameter of 0.88 mm. To quantify the pattern, the fluorescent intensity was measured across a branch (white arrow).

The self-organized patterning can also be superimposed with pre-patterning of seeding cells (Figure 5B). When placing the colonies close by, the colonies first grew independently, forming a single mCherry ring although stretched outwards. Then the cells grew into a connected patch and a mCherry contour formed at the edge of the meta-colony. By day two, a second contour line also formed. This indicates the core-ring dynamic extended beyond simple geometries.

Under certain conditions, the system also demonstrates composite patterns consisting of branches and rings (Figure 5C). This is likely due to the residual (though attenuated) QS capability of PA14-*lasR*. Thus, seeding colonies close-by allows sharing of both AHL and surfactant leading to enhanced branching capability. Following this hypothesis, we generated robust branching-ring hybrid patterns within 22 hours by printing cell culture in close distance (Figure 5D). With longer incubation time, we eventually observed concentric contours in branches. When cells were placed further apart, no branch formed but each colony acted as individuals and formed rings within (Supplementary figure 12). These results indicate the branches and rings formed rather independently. This composability suggests the possibility of “stacking” different patterning capabilities to generate layered patterns.

## Discussion

The fundamental goal of synthetic biology is to engineer cells to generate desired outcomes. Depending on the specific outcomes, this goal can be achieved through high-throughput experiments to screen a sufficiently large library of candidate systems or conditions^34–37^. Alternatively, it can be achieved through directed evolution^38,39^. This approach is most applicable if the target outcome can be generated and evaluated by a relatively simple readout, such as “ON” and “OFF” states (for logic gates)^40^, gene expression levels^34^, growth rates, or enzymatic functions^37^.

For many applications, however, it is difficult to generate or evaluate the target outcomes at high throughput. Examples include oscillations^41,42^, spatial dynamics^5–9,13–15,18^, phenotype plasticity to changing environments^22,43^, and long term evolution^25,44,45^. For these systems, a reliable modeling framework can provide an effective approach to explore system behavior and identify optimal system design or experimental conditions. Our work demonstrates such an approach, which integrates mechanistic modeling and machine learning for fast and interpretable predictions. It is agnostic to specific modeling approaches^13,46–48^ but rather depends on the learnability of dynamics and computational efficiency. In general, it is particularly effective when: 1. high-throughput experimentation is not feasible, 2. a reliable mechanistic model is available but still computationally inefficient.

The key benefit of the ML model in our pipeline is computational acceleration. Though the ML model itself is not interpretable, its predictions can be evaluated by mechanistic modeling, thus enabling interpretability. We note that this interpretability differs from the attempt to interpret the ML model *per se*, which is more challenging. For instance, in our framework, the predictions are made by exploiting a low dimensional representation of the pattern profiles, using latent vectors. These latent vectors will depend on our model architecture and training process, as well as the pattern profiles. As such, direct interpretation of these vectors would be challenging. Despite the difficulty or inability to interpret the latents directly, however, our mechanistic model provides a route to interpret parameter combinations, as well as the rules emerging from the ensemble analysis of many examples. This parallel integration of mechanistic modeling and ML provides a new perspective of providing interpretability when using ML models. While we focused on pattern formation, this approach is generally applicable when dealing with other complex system functions to engineer or optimize.

Our overall computational framework (Figure 2) is modular and generally adaptable for diverse prediction tasks. We used a VAE-MLP architecture to predict the input-output relationships of the patterning system, the architecture is concise and effective in embedding 1D data series with high fidelity, including spatial profile (this study) and time courses^49^. In general, any generative AI models, such as GANs, LSTM^30^, and transformers^50^, or other sophisticated algorithms for solving PDEs^51,52^, can be used for similar tasks^53,54^. When selecting an architecture, key considerations are the model capacity, training process, and the compute speed. In our case, complex pattern formation occupies an extremely small fraction of the parameter space, only a fast enough ML surrogate can generate sufficient representations to derive rules, such as shown in Figure 2E. Compared to our previous efforts using an LSTM^30^, this model gives 100 fold higher acceleration. Additionally, we considered the generation of the training dataset. We envision that with more powerful ML architectures and lighter models, fewer training data and consequently further acceleration can be achieved. Lastly, for pattern formation, we focused solely on the input-output relationships and disregarded the transient dynamics. For systems where temporal dynamics are significant, different models or data representation can be specifically tailored towards this goal. For systems demanding higher spatial accuracy or temporal resolutions, models that take a more integrative approach between the numerical methods and ML may be used^51,52,55,56^. For understanding transient dynamics, methods such as state-space models could be employed^57^. Different models have proven effective for various systems; comprehensive studies on computational efficiency and accuracy improvements should be conducted.

The rules we derived encompass not only circuit dynamics but also environmental factors and host physiology. Indeed, selecting different hosts can open new opportunities to tune system dynamics. Factors such as fitness, motility, tolerance to circuit products, maintenance of the metabolism are often de-emphasized in experiments but can have significant implications. In this endeavor, introducing *E. coli* minD/E patterning system into mammalian hosts has created fascinating intracellular patterns^58^. Host context has been demonstrated as an engineering parameter for programming temporal dynamics^31,32^. For programming metabolic phenotypes, transplanting biosynthetic gene clusters into different prokaryotes and eukaryotes hosts has enabled the discovery of a class of rare microbiome-derived nucleotide metabolites^59^.

Branching *P. aeruginosa* and the synthetic circuit are two composite patterning systems. Composability is a fundamental strategy in nature, observed across all levels – from individual proteins, gene expression systems, metabolic pathways, to cells. Such that multicellular systems can function seamlessly. The key question is to which degree the systems need to be decoupled, and what are the shared properties that link them. If two systems are completely orthogonal, layered patterns can form. If they are partially coupled and the linking properties match within desired ranges, composable patterns can also be designed. For instance, hierarchical induction using two morphogen gradients has been designed to form patterns in different channels^48^. Cell-cell adhesion and juxtacrine signaling systems were used together to create layered, single core or multi core 3D patterns^10^. With increased programmability, it will open doors to leveraging pattern formation in various applications^60–63^.

## Methods

### Host strains and plasmids

PA14 wildtype and *fleN* (V178G) mutant were generous gifts from Xavier lab. The PA14-*lasR* mutant was obtained from spatial evolution^25^ (Genome: NCBI Sequence Read Archive, BioProject PRJNA875393, Strain 36 - SRX17381941). The plasmids were constructed using Gibson assembly. To construct the PA14 gene circuit, PCR was carried out to duplicate the circuit part from the *E.coli* plasmid pET15bLCFPT7, and the circuit fragment was inserted onto the other *E.coli* plasmid pTuLys2CMR2^15^. Next, we inserted pRO1600 replication origin (from pUCP30T-eCFP) and replaced the antibiotic resistance marker by gentamycin resistance gene. The sequence of the final PA14 plasmid (pTu_Lys_T7_GMR_pRO1600) was whole plasmid sequenced by Primordium Labs and confirmed. Other plasmid designs were constructed using similar methods, different shuttle vectors, broad host replication origins, fluorescent proteins, and plasmid backbones were tested.

The plasmid was transformed into PA14 hosts by electroporation^26^. Culture PCR was carried out to confirm the successful transformation. Single colonies of the circuit carrying strains were grown on selective plates and whole genome sequenced (Illumina NovaSeq 6000) by SeqCenter. The samples were prepared with Illumina DNA Prep kit. The reads were aligned to the ensemble *P. aeruginosa* PA14 genome reference (Pseu_aeru_PA14_V1) and the plasmid sequences using *breseq*^64^. Plasmid copy number was estimated by the ratio of mean coverage between the plasmid and chromosome read alignments. The final value was determined by averaging the results from independent sequencing of two single colonies.

### Pattern formation experiments on agar plates

To grow the circuit-carrying *P. aeruginosa* colonies, the cells were inoculated in 3 ml LB medium (Genesee) with 10 μg/ml gentamycin sulfate (VWR) in a culture tube, and grew overnight in a shaker incubator at 37°C and 200 rpm. Before experiments, 200 μl overnight culture was diluted in 1 ml fresh LB medium with gentamycin and incubated at 37℃ with 200 rpm shaking for an additional 3 hours to allow the cells to recover to the exponential growth stage and reach an OD of 0.2 - 0.4 (measured at 600 nm). Before inoculating on agar plates, the cell culture was washed and diluted with LB to an OD of 0.2, unless stated otherwise.

The growth medium recipe was adopted from Luo *et al*.^65^ with modifications. The base medium contains 1X phosphate buffer, 10 g/L casamino acid (Gibco™ Bacto™ 223120), 0.5% agar (w/v, BD Difco™ 214530), 0.1 mM CaCl_2_, 1 mM MgSO_4_, 10 μg/ml gentamycin sulfate, and 1 mM IPTG (GoldBio), unless stated otherwise. The stock solutions were prepared as follows. The stock 5X phosphate buffer was made by mixing 12 g Na_2_HPO_4_ (anhydrous), 15 g KH_2_PO_4_ (anhydrous), and 2.5 g NaCl in deionized water, and sterilized by autoclaving. The stock casamino acid solution was made by dissolving 40 g casamino acid in 200 ml deionized water and sterilized by vacuum filtration. The final concentration was 200 g/L. The stock 1 M MgSO_4_ and 0.1 M CaCl_2_ solutions were made by dissolving the salts in deionized water and sterilized by autoclaving. The agar was melted in sterilized deionized water by microwaving before experiments. The growth medium was made by mixing the stock solutions and was prepared fresh every time.

Regardless of the type of petri dish, we made sure that the agar height was consistent, to remove the impact of agar thickness. 13 ml of the medium was used for 10 cm round petri dishes (Falcon), 15.7 ml was used for one-well plates (Greiner Bio-One). We let the petri dishes solidify at room temperature with the lids closed for 2 hours before inoculation.

To inoculate, 1 μl cell culture was manually pipetted onto the agar surface at the center of the petri dish. The agar plates were left to dry on the bench for 15 min with the lids open next to a Bunsen burner. The plates were then incubated upside down at 37 °C in an incubator without shaking for 24 to 48 hours.

To grow the swarming PA14 colonies in Figure 1C, PA14 wildtype, *fleN* mutant and *lasR* mutant cell cultures were prepared in a similar way but without antibiotics. 20 ml of the standard growth medium, without gentamycin and IPTG, was poured into 10 cm petri dishes. The cells were incubated for 20 hours before imaging.

### Imaging

Fluorescent imaging was done with a Keyence BZ-X710 fluorescence microscope, and Keyence BZX Software Suite 1.3.1.1. A 2X objective was used. mCherry and CFP channels were imaged using RFP and ECFP filter cubes at optimal exposure, or equal exposure for comparison. When colonies grew larger than a single field of view, the entire colony was scanned and stitched together using the built-in algorithm of the microscope. The images were processed with ImageJ. For brightfield images, the intensities were first inverted before quantification. The cross profiles were taken along a line from the colony center to the edge.

### Bioprinting

Bacterial cell culture was prepared in the same way as manual pipetting. Mantis liquid dispenser (FORMULATRIX) was used to print cell culture onto agar surface on a petri dish or one well plate. A low volume microfluidic chip was used to enable printing as little as 0.1 μl. The inoculum diameter depends on the seeding volume and cell density. For OD = 0.2, 0.1 μl gives a diameter of 0.88mm. The diameters for different conditions were measured by imaging and used in modeling. When printing arrays, the configuration was first defined in the Mantis software, which used a template of 396-well microtitre plate for selecting the seeding locations. The minimal spacing is defined by the center-to-center distance between two immediate neighboring wells, which is 2.25 mm. Larger spacings are multiplications of the minimal spacing, e.g. placing seeding positions apart by 1, 2, 3 wells apart leads to 4.5, 6.75, and 9 mm spacing between colony centers.

### Growth measurements

The growth curves were measured using a plate reader (Tecan Infinite 200). In each well of a 96-well plate (Corning), we added 196 μl liquid growth medium (same formula as the solid phase medium except agar), 2 μl OD = 0.02 cell culture, 2 μl 100X IPTG. and 0.2 μl 1000X antibiotics (for plasmid carrying strains). For generating Supplementary figure 3, 0.2 μl 1000X 3OC6HSL (Sigma-Aldrich) stock solution was also supplemented. After adding the solutions, 50 μl mineral oil (Sigma-Aldrich) was added on top to prevent evaporation. The 96-well plates were then incubated in a plate reader at 37°C with shaking every 10 minutes. OD, RFP, and CFP measurements were taken at 10-min intervals for up to 24 hours. Background signals measured from media containing no cells were subtracted from the readouts. The growth rates were calculated from the exponential growth phase OD600 results.

### Mathematical modeling

The model details are described in the Supplementary Information. The model was implemented and solved numerically with MATLAB (R2022a). The numerical simulation was done on Duke Compute Cluster using Intel Xeon Gold 5317 CPU processors at 3.0 GHz. SLURM was used for running parallel jobs on the cluster. The average computation time per data point (Figure 2C) was calculated on a screening job array distributed to multiple CPU cores.

### Machine learning surrogate

The machine learning surrogate model details are described in the Supplementary Information. The ML model was built in PyTorch (Version 2.1). It was trained on Duke Compute Cluster with NVIDIA RTX A1500 GPU, CUDA version 12.2. GPUs were used for large-scale ML screening. However, when benchmarking (Figure 2C), ML inference was done on the same CPU hardware as PDE simulations. The average computation time was calculated by dividing the time for batch inference by the batch size. The batch size was varied, and the converged, average compute time was taken for comparison.

## Code and data availability

The MATLAB and Python code used for PDE simulation, machine learning, and data analysis in the study is available on GitHub: https://github.com/youlab/PatternFormationRules_JiaLu.

## Author contribution

J. L. conceived the research, conducted experiments and computation, and wrote the manuscript. N.L. assisted with gene circuit construction and modeling. K.S. and S.L. carried out experiments. R.M. assisted with sequencing. Y.B. assisted with numerical simulations. L.Y. conceived the research and wrote the manuscript. All authors participated in manuscript revision.

## Acknowledgement

The authors thank S. Wang, E. Simsek for the constructive feedback, J. Xavier group for sharing PA14 strains, L. Diechret group for helpful discussions, and Duke Compute Cluster and T. Milledge for assistance and resources on high performance computing. This work was supported by Office of Naval Research (LY: N00014-12-1-0631) and National Science Foundation (LY: MCB-1937259).

## Conflict of interest

The authors declare no conflict of interest.

## Notes

### Competing Interest Statement

The authors have declared no competing interest.

### Summary of Updates

Title updated; minor corrections were made to the maintext

