## Supplementary information for "Discovery of interpretable patterning rules by integrating mechanistic modeling and deep learning"

### Decoding pattern formation rules by integrating mechanistic modeling and deep learning

**Strains and plasmids**

**Strains**

| | Reference or source | Description/catalog number |
|---|---|---|
| PA14 wildtype | Xavier lab (Memorial Sloan-Kettering Cancer Center, New York) | |
| PA14 *fleN* (V178G) mutant | Xavier lab | |
| PA14 *lasR* mutant | Luo *et al.*[1] | Isolated from evolution |
| DH5α | Thermal Fisher Scientific | EC0112 |
| MG1655 | | |

**Plasmid**

| | Reference or source | Description |
|---|---|---|
| pET15bLCFPT7 | Cao *et al.*[2] | Positive feedback part of the circuit for *E.coli* |
| pTuLys2CMR2 | Cao *et al.*[2] | Negative feedback part of the circuit for *E.coli* |
| pUCP30T-eCFP | Addgene | pRO1600 doner |
| pTu_Lys_T7_GMR_pRO1600 | | The full patterning circuit for *P. aeruginosa*, including *lacI* |
| mCh_Lys_GMR_pRO1600 | | Negative feedback part of the circuit for *P. aeruginosa*. Containing plux promoter, lysozyme, pT7 promoter, *mCherry*, *luxI*, *luxR* genes. Carrying pRO1600 fragment and gentamycin resistance gene |
| pUCP30T_T7_GMR_lacI_pRO1600 | | Positive feedback part of the circuit for *P. aeruginosa*. Containing pT7 promoter, |

| | |
|---|---|
| | *T7RNAP*, *CFP*, *lacI* genes. Carrying pRO1600 fragment and gentamycin resistance gene. |
| pUCP30T_T7_lys_lasR_GMR_pRO1600 | The full patterning circuit for *P. aeruginosa*, *luxR* was replaced by *lasR* gene. It carries pT7 promoter, *T7RNAP*, *CFP*, *luxI*, *lasR*, plux promoter, *T7 lysozyme*, *mCherry*, and *lacI* genes. It also has pRO1600 fragment and gentamycin resistance gene. |

#### Reagents and tools

##### Chemicals, enzymes, and reagents

| Reagent/Resource | Reference or source | Catalog number/ version |
|---|---|---|
| Na2HPO4 (anhydrous) | Sigma-Aldrich | S3264 |
| KH2PO4 (anhydrous) | Sigma-Aldrich | P5655 |
| NaCl | Sigma-Aldrich | S3014 |
| MgSO4 | Sigma-Aldrich | M7506 |
| CaCl2 | Sigma-Aldrich | C1016 |
| Casamino acids | BD | Bacto 223120 |
| Granulated agar | BD | Difco 214530 |
| Gentamycin sulphate | VWR | 97061 |
| 3OC6HSL | Sigma-Aldrich | K3007 |
| 3OC12HSL | Sigma-Aldrich | O9139 |
| IPTG | GoldBio | I2481C |

##### Software

| Reagent/Resource | Reference or source | Version |
|---|---|---|
| MATLAB | The MathWorks Inc. | R2022a |
| Python | | 3.9 |
| PyTorch | Paszke *et al.*[3] | 2.1 |
| breseq | Deatherage and Barrick[4] | |
| Keyence BZX Software Suite | Keyence | 1.3.1.1. |

| | | |
|---|---|---|
| ImageJ | Schneider *et al.*[5] | 1.53k |
| Doc-It Colony Counter software | Analytik Jena | 7.0.3 |
| PyMol | Schrödinger | 2.5.4 |

**Equipment**

| Reagent/Resource | Reference or source | Version |
|---|---|---|
| MANTIS automated liquid handler | FORMULATRIX | |
| UVP Colony Doc-It Imaging Station | Analytik Jena | |
| Plate reader | Tecan | Infinite 200 |
| Keyence microscope | Keyence | BZ-X710 |

### 1. Mathematical modeling

#### 1.1. Model formulation

The mechanistic model was developed to describe the colony development and pattern formation of the PA14-*lasR* carrying the patterning circuit. The model accounts for cell proliferation, cell motility, circuit gene expression and interaction, AHL synthesis and diffusion, nutrient consumption and diffusion. It was modified based on Cao *et al* [6]. In the following equations, C, N, A, B, T, L, and P correspond to cell density, nutrient concentration, 3OC6HSL (product of LuxI), 3OC12HSL (product of LasI), T7 RNA polymerase (T7RNAP), T7 lysozyme, and T7RNAP-lysozyme complex, respectively. T, L, and P represent averaged intracellular protein concentrations.

The cell motility is approximated by a diffusion term. Cell growth is modeled after logistic growth (Monod function) and is subject to burden from circuit expression. The Monod function accounts for the contribution of nutrient to the overall colony growth, while the logistic term accounts for the limit of cell growth at a specific location. As the colony is small (<5mm) compared to the entire growth domain (e.g. a 10cm petri dish) and nutrient is sufficiently supplemented, the cell carrying capacity is unlikely to be limited by nutrient availability. It is also assumed that the nutrient can diffuse laterally in agar.

The gene circuit expresses 3OC6HSL, a quorum sensing small molecule produced by LuxI. It can diffuse in agar. Its production is governed by the upstream T7 promoter activity (driving LuxR and LuxI expression), which is regulated by T7 RNA polymerase and T7-lysozyme complex. The host stain PA14-*lasR* has an intact *lasI* gene (encoded on chromosome) which leads to the synthesis of another quorum sensing small molecule 3OC12HSL, as a part of the host's quorum sensing systems. Its expression is incorporated as a function of cell density and gene expression profile. This molecule also mediates host-circuit crosstalk, by binding with luxR protein to upregulate pLux promoter activity. Thus, the Hill equations of A and B are added up to for the pLux promoter activity.

T7 RNA polymerase and T7 lysozyme are intracellular proteins, and they can form a complex inhibiting the activity of pT7 promoter. These two proteins' expression is regulated by pT7 promoter and pLux promoter, respectively. T, L, and P are the local protein concentrations per cell. Thus, as the cells move, their values change accordingly, described by the passive tracer terms. The derivation can be found in Cao *et al*[2].

The PA14-*lasR* exhibits weakened swarming capability, typically forming a small singular colony. When inoculated at higher initial cell densities or inoculated at multiple neighboring seeding positions, this strain can still form branching colonies. The PA14-*lasR* patterning system forms a small circular colony in the patterning experiments, thus it is viewed as radial symmetric, captured by this model. Branching mechanism is not implemented here.

There are several key differences between this model and the one in Cao *et al*[2]. In our experiments, cells grew on agar petri dish without any vertical confinement, and the petri dishes were significantly larger than the colony size. For example, a petri dish has a diameter of 10 cm, and a one-well plate is sized at 128 mm × 86 mm, whereas the colony diameter is less than 5mm. On the other hand, Cao *et al.* grew *E. coli* colonies on a soft, thin agar pad on an imaging slide.

The colonies were clamped by a cover glass from top confining the colony height. Thus, we removed the colony height confinement and allowed for free diffusion of nutrient and signaling molecules, instead of assuming even distributions. Besides, PA14-*lasR* has intact *lasI* gene encoded on the chromosome. Thus, 3OC12HSL, the product of *lasI* is introduced in our model. Its synthesis, diffusion, and crosstalk with the circuit are incorporated.

The model is formulated as follows:

$$\frac{\partial C}{\partial t} = D_C\nabla^2 C + \alpha_C \frac{1}{1+\alpha T+\beta L}\frac{N}{N+K_N}\left(1-\frac{C}{\bar{\bar{C}}}\right)C$$

$$\frac{\partial N}{\partial t} = D_N\nabla^2 N - \beta_N\alpha_C \frac{1}{1+\alpha T+\beta L}\frac{N}{N+K_N}\left(1-\frac{C}{\bar{\bar{C}}}\right)C$$

$$\frac{\partial A}{\partial t} = D_A\nabla^2 A + \alpha_A \frac{K_P}{P+K_P}\frac{T}{T+K_T}C\phi(x,C) - d_A A$$

$$\frac{\partial B}{\partial t} = D_B\nabla^2 B + \alpha_B C\phi(x,C) - d_B B$$

$$\frac{\partial T}{\partial t} = D_C\frac{\nabla T\cdot\nabla C}{C} - T\alpha_C \frac{1}{1+\alpha T+\beta L}\frac{N}{N+K_N}\left(1-\frac{C}{\bar{\bar{C}}}\right) - d_T T + \alpha_T\frac{T}{T+K_T}\frac{K_P}{P+K_P}\phi(x,C) - k_1 TL + k_2 P$$

$$\frac{\partial L}{\partial t} = D_C\frac{\nabla L\cdot\nabla C}{C} - L\alpha_C \frac{1}{1+\alpha T+\beta L}\frac{N}{N+K_N}\left(1-\frac{C}{\bar{\bar{C}}}\right) - d_L L + \alpha_L\frac{T}{T+K_T}\left(\frac{A^m}{A^m+K_A^m}+\frac{B^n}{B^n+K_B^n}\right)\phi(x,C) - k_1 TL + k_2 P$$

$$\frac{\partial P}{\partial t} = D_C\frac{\nabla P\cdot\nabla C}{C} - P\alpha_C \frac{1}{1+\alpha T+\beta L}\frac{N}{N+K_N}\left(1-\frac{C}{\bar{\bar{C}}}\right) + k_1 TL - k_2 P$$

$\phi(x, C)$ is the gene expression profile, which gradually increases and saturates towards the colony edge. It is formulated as

$$\phi(\mathrm{x,C}) = \begin{cases} \dfrac{K_\phi^l}{K_\phi^l + (R_\phi - x)^l}, & x \le R_\phi \\ 1, & x > R_\phi \end{cases}$$

, where $R_\phi$ is the distance at which the cell density reaches 90% of its maximum value. As the colony is relatively thin, vertical gene expression profile is neglected. The base values of the parameters are summarized in Table 1.

#### 1.2. Non-dimensionalization

To derive the non-dimensional model, we assume the following state variables,

$$\hat{t} = \alpha_C t, \qquad \hat{x} = \frac{1}{\mathcal{L}}x, \qquad \hat{C} = \frac{C}{\bar{C}}, \qquad \hat{N} = \frac{N}{N_0}$$

$$\hat{A} = \frac{A}{K_A}, \qquad \hat{\mathrm{B}} = \frac{B}{K_B}, \qquad \hat{L} = \frac{d_L}{\alpha_L}L, \qquad \hat{T} = \frac{d_T}{\alpha_T}T, \qquad \hat{P} = \frac{P}{K_P}$$

, where they are rescaled by scaling factors.

Plugging these into the model, we get the non-dimensionalized model as follows:

$$\frac{\partial \hat{C}}{\partial \hat{t}} = G_1 \nabla^2 \hat{C} + \frac{1}{1 + \alpha' \hat{T} + \beta' \hat{L}} \frac{\hat{N}}{\hat{N} + G_2} (1 - \hat{C}) \hat{C}$$

$$\frac{\partial \hat{N}}{\partial \hat{t}} = G_3 \nabla^2 \hat{N} - G_4 \frac{1}{1 + \alpha' \hat{T} + \beta' \hat{L}} \frac{\hat{N}}{\hat{N} + G_2} (1 - \hat{C}) \hat{C}$$

$$\frac{\partial \hat{A}}{\partial \hat{t}} = G_5 \nabla^2 \hat{A} + G_6 \frac{\hat{T}}{\hat{T} + G_7} \frac{1}{\hat{P} + 1} \hat{C} \phi(\hat{x}, \hat{C}) - G_8 \hat{A}$$

$$\frac{\partial \hat{\mathrm{B}}}{\partial \hat{t}} = G_9 \nabla^2 \hat{\mathrm{B}} + G_{10} \hat{C} \phi(\hat{x}, \hat{C}) - G_{11} \hat{\mathrm{B}}$$

$$\frac{\partial \hat{T}}{\partial \hat{t}} = G_1 \frac{\nabla \hat{T} \cdot \nabla \hat{C}}{\hat{C}} - \hat{T} \frac{1}{1 + \alpha' \hat{T} + \beta' \hat{L}} \frac{\hat{N}}{\hat{N} + G_2} \left(1 - \hat{C}\right) - G_{12} \hat{T} + G_{12} \frac{\hat{T}}{\hat{T} + G_7} \frac{1}{\hat{P} + 1} \phi\left(\hat{x}, \hat{C}\right)$$

$$- G_{13} \hat{L} \hat{T} + G_{14} \hat{P}$$

$$\frac{\partial \hat{L}}{\partial \hat{t}} = G_1 \frac{\nabla \hat{L} \cdot \nabla \hat{C}}{\hat{C}} - \hat{L} \frac{1}{1 + \alpha' \hat{T} + \beta' \hat{L}} \frac{\hat{N}}{\hat{N} + G_2} \left(1 - \hat{C}\right) - G_{15} \hat{L}$$

$$+ G_{15} \frac{\hat{T}}{\hat{T} + G_7} \left( \frac{\hat{A}^m}{\hat{A}^m + 1} + \frac{\hat{B}^n}{\hat{B}^n + 1} \right) \phi\left(\hat{x}, \hat{C}\right) - G_{16} \hat{L} \hat{T} + G_{17} \hat{P}$$

$$\frac{\partial \hat{P}}{\partial \hat{t}} = G_1 \frac{\nabla \hat{P} \cdot \nabla \hat{C}}{\hat{C}} - \hat{P} \frac{1}{1 + \alpha' \hat{T} + \beta' \hat{L}} \frac{\hat{N}}{\hat{N} + G_2} \left(1 - \hat{C}\right) + G_{18} \hat{L} \hat{T} - G_{19} \hat{P}$$

, the non-dimensional parameters are summarized in Table 2.

mCherry ($\widehat{\Psi_{mCh}}$) and CFP ($\widehat{\Psi_{CFP}}$) co-express with lysozyme and T7RNAP, respectively. Considering that they have leaky expression, small basal expression rates are included for both fluorescent proteins ($a$, $b$). Their nondimensionalized averaged intracellular concentrations are:

$$\frac{\partial \widehat{\Psi_{CFP}}}{\partial \hat{t}} = G_1 \frac{\nabla \widehat{\Psi_{CFP}} \cdot \nabla \hat{C}}{\hat{C}} - \widehat{\Psi_{CFP}} \frac{1}{1 + \alpha' \hat{T} + \beta' \hat{L}} \frac{\hat{N}}{\hat{N} + G_2} \left(1 - \hat{C}\right) + G_{12} \frac{\hat{T}}{\hat{T} + G_7} \frac{1}{\hat{P} + 1} \phi\left(\hat{x}, \ \hat{C}\right) + \ a$$

$$\frac{\partial \widehat{\Psi_{mCh}}}{\partial \hat{t}} = G_1 \frac{\nabla \widehat{\Psi_{mCh}} \cdot \nabla \hat{C}}{\hat{C}} - \widehat{\Psi_{mCh}} \frac{1}{1 + \alpha' \hat{T} + \beta' \hat{L}} \frac{\hat{N}}{\hat{N} + G_2} \left(1 - \hat{C}\right) + G_{15} \frac{\hat{T}}{\hat{T} + G_7} \left( \frac{\hat{A}^m}{\hat{A}^m + 1} + \frac{\hat{B}^n}{\hat{B}^n + 1} \right) \phi\left(\hat{x}, \ \hat{C}\right) + b$$

The accumulative concentrations of protein (e.g. T, L, P, $\Psi_{mCh}$, and $\Psi_{CFP}$) are calculated as

$$W = \widehat{W_{intra}} \cdot \hat{C}$$

, where $\widehat{W_{intra}}$ is the averaged intracellular concentration of the protein. The mCherry pattern used in this work is the accumulative mCherry concentration.

##### 1.3. Boundary conditions

No-flux boundary condition was chosen because of the experimental setup. The petri dish is much larger than the colony size and the cells and chemicals are unlikely to diffuse to the edge of the petri dish within the time period of the experiments given their diffusivity. Thus, to save computational resources and time, we chose a relatively large domain size in the numerical simulation, instead of simulating the entire petri dish, and terminated the simulation before cells and molecules diffused beyond the boundary.

##### 1.4. Initial conditions

In experiments, the cell culture was pipetted onto agar surface and followed by air drying, forming a ring-shaped deposition. We modeled the initial cell distribution after this observation. Besides, seeding volume and cell density impact the inoculum diameter and cell distribution. Thus, they were incorporated based on imaging measurement. For the analysis in Figure 3, the default cell density was 0.2, the default seeding volume was 0.1 μl, which is the base case. This represents the experimental base case of OD = 0.2, seeding volume = 0.1 μl. For analysis in Figure 4, when screening seeding volumes, the cell density was fixed to 0.2, and the diameters corresponding to 0.1, 0.2, 0.3, 0.4, and 1 μl were scaled to 1, 1.3, 1.68, 1.81, and 2.13, respectively, relative to the base case. Similarly, when screening seeding cell density, the seeding volume was fixed to 0.1 μl, and the diameters corresponding to cell density = 0.02, 0.2, and 1 were scaled to 1.15, 1, and 0.85, respectively, relative to the base case. The shape of the cell distribution was determined by the cell number, i.e. a higher total cell number results in a taller $C0$. The shape was scaled with respect to the base case as well.

At $t = 0$, nutrient was uniformly distributed, characterized by the initial concentration $N_0$. 3OC6HSL and 3OC12HSL started with zero concentrations as the cells have been washed, and they have not yet produced any LuxI and LasI. Lysozyme and the T7RNAP-lysozyme complex were also assigned zero initial concentrations. T7 RNA polymerase was assumed to have a small amount of leaky expression. It was assumed to start with a marginal concentration to initiate the circuit transcription, proportional to the initial cell density.

##### 1.5. Finite difference method

The patterns and colonies are radially symmetric. Thus, the model can be numerically solved in one dimension. Polar coordinate was used, and the only spatial variable was the distance to colony center, namely $r \in [0, R]$. The equations were split into three components – advection, diffusion, and reaction. At each time step $\Delta t$, each part was solved sequentially. First order implicit time scheme and first order central difference spatial scheme were used to solve the advection and diffusion terms, and MATLAB's built-in ODE45 was used to solve the reaction terms. To ensure the numerical methods are robust, we conducted convergence analysis, where different grid size and time step were tested. For each grid size $h$ and time step $\Delta t$, 9 sets of numerical simulations were generated from different PDE parameter combinations. To calculate the error, the numerical results generated from the smallest $\Delta t$ and the finest $h$ were used as the norm, based on which $R^2$ was calculated. The error was fitted to the power law. The finite difference method is indeed convergent, with a convergence rate ~1.5. Based on the analysis, $\Delta t$ = 0.025 and $h = 0.01$ were chosen as the default setup. The total computational time was set to 20.

##### 1.6. Parameter Tables

**Table 1. Dimensional parameters.**

| | Parameters | Description | Base values | Units | References |
|---|---|---|---|---|---|
| 1 | $D_C$ | Diffusion coefficient of PA14 *lasR* mutant on swarming agar plates | 2.5E-3 | $cm^2 \cdot h^{-1}$ | Song *et al.*, 2009[7] |
| 2 | $D_N$ | Diffusion coefficient of nutrient | 0.162 | $cm^2 \cdot h^{-1}$ | Luo, Lu, Simsek *et al.*, 2024[1] |
| 3 | $D_A$ | Diffusion coefficient of 3OC6HSL | 0.06 | $cm^2 \cdot h^{-1}$ | Ravichandar *et al.*, 2017 [8] |
| 4 | $D_B$ | Diffusion coefficient of 3OC12HSL | 0.06 | $cm^2 \cdot h^{-1}$ | Ravichandar *et al.*, 2017 [8] |
| 5 | $\alpha_C$ | Cell growth rate | 0.2 | $h^{-1}$ | Estimated based on experiments |
| 6 | $\alpha_A$ | Synthesis rate of 3OC6HSL by *luxI* | 1000 | $nM \cdot h^{-1}$ | You *et al.*, 2004[9] |
| 7 | $\alpha_B$ | Synthesis rate of 3OC12HSL by *lasI* | 100 | $nM \cdot h^{-1}$ | Estimated based on experiments |
| 8 | $\alpha_T$ | Synthesis rate of T7RNAP | 800 | $nM \cdot h^{-1}$ | Cao *et al.*, 2016[2] |
| 9 | $\alpha_L$ | Synthesis rate of lysozyme | 500 | $nM \cdot h^{-1}$ | Cao *et al.*, 2016[2] |
| 10 | $\beta_N$ | Nutrient consumption rate | 155 | $nM \cdot h^{-1}$ | Cao *et al.*, 2016[2] |
| 11 | $d_A$ | Degradation rate of 3OC6HSL | 0.1 | $h^{-1}$ | You *et al.*, 2004[9] |
| 12 | $d_B$ | Degradation rate of 3OC12HSL | 0.1 | $h^{-1}$ | Balagaddé *et al.*, 2008[10] |

| | | | | | |
|---|---|---|---|---|---|
| 13 | $d_T$ | Degradation rate of T7RNAP | 0.3 | $h^{-1}$ | Payne *et al.*, 2013[11] |
| 14 | $d_L$ | Degradation rate of lysozyme | 0.0144 | $h^{-1}$ | Payne *et al.*, 2013[11] |
| 15 | $k_1$ | Association rate of T7RNAP and lysozyme | 400 | $nM\cdot h^{-1}$ | Kumar and Patel, 1997[12] |
| 16 | $k_2$ | Disassociation rate of T7RNAP-lysozyme complex | 10,800 | $h^{-1}$ | Kumar and Patel, 1997[12] |
| 17 | $K_N$ | Half-saturation for nutrient uptake | 20,000 | nM | Estimated based on experiments |
| 18 | $K_P$ | Half inhibition of T7RNAP - lysozyme complex | 400 | nM | Cao *et al.*, 2016[2] |
| 19 | $K_T$ | Half activation constant of T7RNAP | 1200 | nM | Cao *et al.*, 2016[2] |
| 20 | $K_A$ | Concentration threshold of 3OC6HSL to half maximum of the pLuxI promoter | 20 | nM | Collins *et al.*, 2006[13] |
| 21 | $K_B$ | Concentration threshold of 3OC12HSL binding with luxR to half maximum of the pLuxI promoter | 1000 | nM | Estimated based on Wu *et al.*, 2014 [14] |
| 22 | $K_\varphi$ | Half activation distance for gene expression | 0.3 | cm | Cao *et al.*, 2016[2] |
| 23 | $\alpha$ | Inhibition factor of T7RNAP on cell growth | 1 | | Cao *et al.*, 2016[2] |
| 24 | $\beta$ | Inhibition factor of T7 lysozyme on cell growth | 100 | | Cao *et al.*, 2016[2] |
| 25 | $\bar{C}$ | Cell carrying capacity | $5 \times 10^3$ | $mm^{-2}$ | Cao *et al.*, 2016[2] |
| 26 | $m$ | Hill coefficient of *luxI* mediated gene expression | 2 | | Payne *et al.*, 2013[11] |

| | | | | | |
|---|---|---|---|---|---|
| 27 | $n$ | Hill coefficient of *lasI* mediated gene expression | 2 | | Wu *et al.*, 2014[14] |
| 28 | $l$ | Hill coefficient for distance-dependent gene expression capacity | 1 | | Cao *et al.*, 2016[2] |
| 29 | $a$ | Basal CFP expression rate | 5 | $nM \cdot h^{-1}$ | Estimated based on Wu *et al.*, 2014[14] |
| 30 | $b$ | Basal mCherry expression rate | 5 | $nM \cdot h^{-1}$ | Estimated based on Wu *et al.*, 2014[14] |
| 31 | $N_0$ | Initial nutrient concentration | 30,000 | nM | Estimated based on experiments |

**Table 2. Dimensionless parameters.**

| | **Parameter** | **Description** |
|---|---|---|
| 1 | G1 | $\frac{D_C}{\alpha_C \mathcal{L}^2}$ |
| 2 | G2 | $\frac{K_N}{N_0}$ |
| 3 | G3 | $\frac{D_N}{\alpha_C \mathcal{L}^2}$ |
| 4 | G4 | $\frac{\beta_N \bar{C}}{N_0}$ |
| 5 | G5 | $\frac{D_A}{\alpha_C \mathcal{L}^2}$ |
| 6 | G6 | $\frac{\alpha_A \bar{C}}{\alpha_C K_A}$ |
| 7 | G7 | $\frac{k_T d_T}{\alpha_T}$ |
| 8 | G8 | $\frac{d_A}{\alpha_C}$ |
| 9 | G9 | $\frac{D_B}{\alpha_C \mathcal{L}^2}$ |

| | | |
|---|---|---|
| 10 | G10 | $\frac{\alpha_B \bar{C}}{\alpha_C K_B}$ |
| 11 | G11 | $\frac{d_B}{\alpha_C}$ |
| 12 | G12 | $\frac{d_T}{\alpha_C}$ |
| 13 | G13 | $\frac{k_1 \alpha_L}{\alpha_C d_L}$ |
| 14 | G14 | $\frac{k_2 K_P d_T}{\alpha_C \alpha_T}$ |
| 15 | G15 | $\frac{d_L}{\alpha_C}$ |
| 16 | G16 | $\frac{k_1 \alpha_\mathrm{T}}{\alpha_\mathrm{C} d_\mathrm{T}}$ |
| 17 | G17 | $\frac{k_2 K_P d_L}{\alpha_C \alpha_L}$ |
| 18 | G18 | $\frac{k_1 \alpha_\mathrm{T} \alpha_L}{K_P \alpha_\mathrm{C} d_\mathrm{T} d_L}$ |
| 19 | G19 | $\frac{k_2}{\alpha_C}$ |
| 20 | $\alpha'$ | $\frac{\alpha \alpha_T}{d_T}$ |
| 21 | $\beta'$ | $\frac{\beta \alpha_L}{d_L}$ |

#### 2. Machine learning surrogate model

##### 2.1. Model architecture and training

A multilayer perceptron-variational autoencoder (MLP-VAE) architecture was used for the machine learning (ML) surrogate model. As the name suggests, the model consists of two parts: an MLP and a VAE decoder. VAE is a generative model that can learn to compress 1D patterns into lower-dimensional vectors (encoder), which are called latent variables, and then decompresses them back into the original higher-dimensional patterns (decoder). Additionally, there is an MLP that takes the 12 PDE parameters as inputs and predicts the VAE latent variables. The surrogate model is the MLP connected to the VAE decoder, in this way, it emulates a PDE simulation — taking PDE parameters as inputs and predicting the simulated pattern.

To efficiently handle 1D patterns, the VAE encoder and decoder are composed of 1D convolutional and deconvolutional layers, respectively. The MLP uses fully connected layers.

The training was done in two steps. First, we trained a VAE using simulated 1D mCherry profiles. Kaiming uniform initialization was used to initialize the weights. Adam optimizer, learning rate scheduling (e.g. ExponentialLR and LambdaLR from PyTorch) and clamping, warm-up, and early stopping were used for consistent and efficient training progression. The reconstruction loss is the sum of mean squared error loss (MSE) and Kullback–Leibler (KL) divergence, where the KL divergence is scaled by a factor $\alpha = 2\times10^{-5}$.

Next, we trained a MLP to map the PDE parameters into VAE latent variables. Instead of updating MLP weights by comparing the latents, we let the predicted latents to pass through the trained VAE decoder and examined the predicted 1D profiles. In other words, the MLP was connected to the trained VAE decoder, which was frozen during MLP training. The same reconstruction loss was used here for updating MLP weights. The training data was the PDE parameter combinations and the corresponding 1D patterns. Adam optimizer, learning rate scheduling and clamping, warm-up, and early stopping were again used for consistent and efficient training progression.

For inference, the trained MLP and trained VAE decoder were connected, such that a PDE parameter combination was given to the surrogate model to predict the mCherry cross profile.

To evaluate model performance, $R^2$ score was reported on train and test sets for both training steps. Let $y_i$, $\hat{y_i}$ be the true and predicted pattern profile, respectively. $R^2$ is defined as $R^2 = 1 - \frac{SS_{res}}{SS_{tot}}$ , where $SS_{res} = \sum_i (y_i - \hat{y_i})^2$ , $SS_{tot} = \sum_i (y_i - \bar{y})^2$. Higher value indicates higher accuracy.

The latent dimension is a hyperparameter shared by both VAE and MLP and impacts model capacity. To finalize model architecture, different latent dimensions were screened, and 16 was chosen for the final model, as it gave the highest accuracy. The final model architecture is as follows. The MLP consists of 6 fully connected (FC) layers, each followed by a ReLU activation function. The model input dimension is 12, and the FC layer dimension increases to as wide as 128 and then decreases throughout the layers to 32. The outputs from the last layer are passed to two separate FC layers, which are responsible for producing the VAE mean and log variance (logvar), respectively. Their dimension (latent dimension) is 16. The VAE encoder is composed of six 1D convolutional layers, and outputs a mean and a log variance, which are then fed into a reparameterization module. The reparameterization module converts them into a single latent vector, z. This vector is subsequently fed into the decoder, which is made of six 1D deconvolutional layers. The model input and output both have dimensions of 201, corresponding to the dimension of the 1D profile. When connecting the MLP to the VAE, the MLP outputs are fed into a reparameterization module, which converts the mean and log variance into the latent vector z in the same way as the VAE counterpart. This latent vector z is then passed into the VAE decoder.

##### 2.2. Training data

The training dataset is made of 30,000 numerical simulation data points. To construct it, 12 partial differential equation (PDE) parameters were selected for parameter screening based on two criteria: their potential significant impact on pattern formation and their experimental tunability. Their screening ranges and methods (either logarithmic or linear) were determined according to their biological dynamical ranges and meanings (Supplementary Table 1). A total of 30,000 numerical simulations were done, each using a randomly sampled combination of these parameters.

During data preprocessing prior to training the ML surrogate, the simulated mCherry profile was first downsampled from dimension 1001 to 201 by keeping one value for every five values. The pattern was then normalized with respect to its maximum value. The PDE parameters were normalized according to their respective ranges and sampling scales and rounded to 8 digits, thus they all ranged from 0 to 1. The normalized PDE parameter combinations and normalized mCherry profiles were used as training data. 10% of the data was used as the test set, 10% as the validation set and the rest as the training set.

Training dataset size impacts ML model accuracy. More data generally results in higher accuracy but would require longer simulation time. To balance the two factors, we screened different dataset sizes. Remarkably, even with a training dataset as small as ~7,000 data points, the model achieved an $R^2 > 0.96$ on test set. For the final model, we trained using 26,526 training data to ensure the best possible performance.

##### 2.3. Pattern characterization

Regardless of whether the pattern was simulated or predicted by the ML model, the number of rings was determined by counting the number of peaks in the full mCherry profile using the same algorithm. The pattern was first flipped and concatenated to construct the full 1D cross profile, extending from one end to the other across the colony center. The number of peaks was then identified using SciPy (v1.11)'s find_peaks function, which recognizes peaks by finding all local maxima through comparing neighboring values. Features such as the distance between maxima, peak width, peak height, and prominence were adjusted to ensure that significant peaks are correctly identified. We categorized 1–2 peaks as 1 ring, 3–4 peaks as 2 rings, 5–6 peaks as 3 rings, and so on. We aimed to identify conditions that produce two or more rings, as these patterns are rare or have never been observed in *E. coli* systems.

##### 2.4. Computational validation

During computational validation, we randomly generated 16 millions of the 12-parameter combinations between 0 and 1, rounded to 8 digits. These combinations were then fed into the trained ML surrogate model to generate normalized mCherry profiles, which were categorized into 1, 2, or 3 rings. From each category, 5,000 data points were randomly selected. The corresponding parameter combinations were converted back into their actual ranges using the same ranges that were used for generating the PDE simulations. The rescaled parameters were then input into the PDE simulation, which was processed in the same way as the training data preprocessing.

Finally, the ML predicted profiles and the normalized PDE outputs were compared. The profiles with matching ring numbers, or $R^2 > 0.8$, were considered to have been validated as accurate. Out of all 15,000 samples, 56.06 % were validated as correct.

To construct the dataset for rule derivation, 2,000 samples were randomly taken from the validated 1-ring category, 1,000 samples from the 2-ring category, and 1,000 samples from the 3-ring category, resulting in a validated dataset of 4,000 samples. The overall $R^2$ of this dataset against the ML prediction is 0.8317, as shown in Figure 3B.

During interpolation inference by ML, a small fraction of ML predictions can be slightly noisy. For these data, moving average filter was applied to remove the small noises before identifying the pattern type. Besides, due to the constrain of boundary condition, mCherry signal cannot diffuse beyond the boundary. If $d_N > 0.05 \times \max(d_1, d_2, \ldots, d_N)$, the sample was discarded.

##### 2.5. Data augmentation and model fine-tuning

With the distributional training set, the ML model performance was already outstanding. A small caveat is that the $R^2$ declines for increasing ring numbers. We wondered if this was due to the extremely low representation of multiple ring examples in the original training set (Figure 2E).

Thus, we augmented the dataset by including more 2 and 3 ring samples. Specifically, the parameter combinations generating more than 1 ring was identified in the original dataset. A small random perturbation was added to them such that it creates a new neighboring parameter combination. Then it was given to the PDE simulation to solve for the 1D profiles. Through this method, we generated 25, 880 samples of two and more rings, and added to the original dataset.

We fine-tuned the ML surrogate model using the augmented dataset, which only slightly improved $R^2$ on each of the rarer pattern classes. $R^2$ on 2 rings improved from 0.9364 to 0.9461; $R^2$ on 3 rings improved from 0.7574 to 0.7636. $R^2$ on 1 ring remained almost the same, changing from 0.9778 to 0.9736. Overall, high fidelity quantitative prediction on multiple rings can be challenging. A possible explanation is that these patterns are very sensitive to the input parameters, due to the intrinsic system dynamics. Performance may be improved by incorporating a more sophisticated method combining numerical method and machine learning, alternative to predicting the input-output relationship. However, this is not essential for our purpose of generating coarse-grained rules, as these rules are based on randomly sampled and then validated parameter combinations.

**Supplementary tables**

**Supplementary Table 1.** Key parameters selected for constructing ML training dataset.

12 PDE parameters were chosen due to their potential impact on pattern formation and their experimental tunability. $D_C$ can be tuned by changing agar density. $\alpha_C$ can be tuned by changing nutrient concentration. $\alpha_T$ is regulated by IPTG induction, thus tuned by IPTG concentration. $\alpha_L$ is controlled by pLux promoter, thus tunable by AHL. $d_A$ and $d_B$ can be tuned through pH. Due to the structural similarity between 3OC6HSL and 3OC12HSL, we assume these two parameters are equal and regulated simultaneously. $d_T$ and $d_L$ can be changed by adding protein degradation tags. α and β indicate the metabolic burden caused by expressing T7RNAP and lysozyme. $K_\varphi$ defines the gene expression profile, which vary based on environmental conditions and host strains. $N_0$ is the initial nutrient concentration, part of the growth medium. These 12 parameters were randomly sampled to construct the ML training dataset, while the rest of the parameters were fixed at the base values. The sampling scales were determined by the parameter's biological meaning and experiments.

| | **Parameter** | **Screening ranges** | **Sampling scale** |
|---|---|---|---|
| 1 | $D_C$ | 0.5E-3 – 12.5E-3 | Logarithmic |
| 2 | $\alpha_C$ | 0.15 – 0.5 | Linear |
| 3 | $\alpha_A$ | 500 – 10000 | Logarithmic |
| 4 | $\alpha_T$ | 80 – 8000 | Logarithmic |

| | | | |
|---|---|---|---|
| 5 | $\alpha_L$ | 50 - 5000 | Logarithmic |
| 6 | $d_A\ (= d_B)$ | 0.01 – 1 | Logarithmic |
| 7 | $d_T$ | 3 – 300 | Logarithmic |
| 8 | $d_L$ | 0.144 – 14.4 | Logarithmic |
| 9 | $\alpha$ | 1 – 5 | Linear |
| 10 | $\beta$ | 2 – 2000 | Logarithmic |
| 11 | $K_\varphi$ | 0.1 – 0.6 | Linear |
| 12 | $N_0$ | 2000 – 50000 | Logarithmic |

#### Supplementary figures

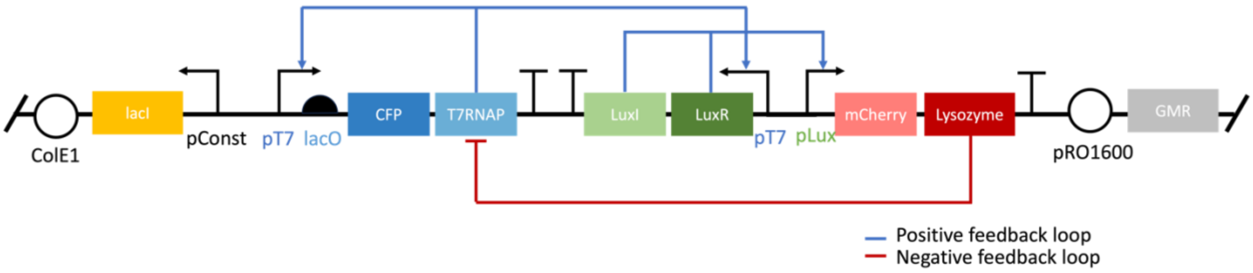


**Supplementary Figure 1.** Diagram of the patterning circuit. In the circuit, T7 RNA polymerase (T7RNAP) serves as the activator by activating its own expression and that of T7 lysozyme. T7 lysozyme is the inhibitor which binds with T7RNAP forming a complex and inhibits T7RNAP expression through repressing the upstream T7 promoter. 3OC6HSL is a quorum sensing (QS) small molecule (produced by LuxI) that freely diffuses in agar and in colony, connecting the activator and inhibitor nodes of the circuit. pRO1600 is a replication origin compatible with *P. aeruginosa*. ColE1 is a replication origin for cloning in *E. coli*. Gentamycin resistance gene (GMR) is used as selection marker.

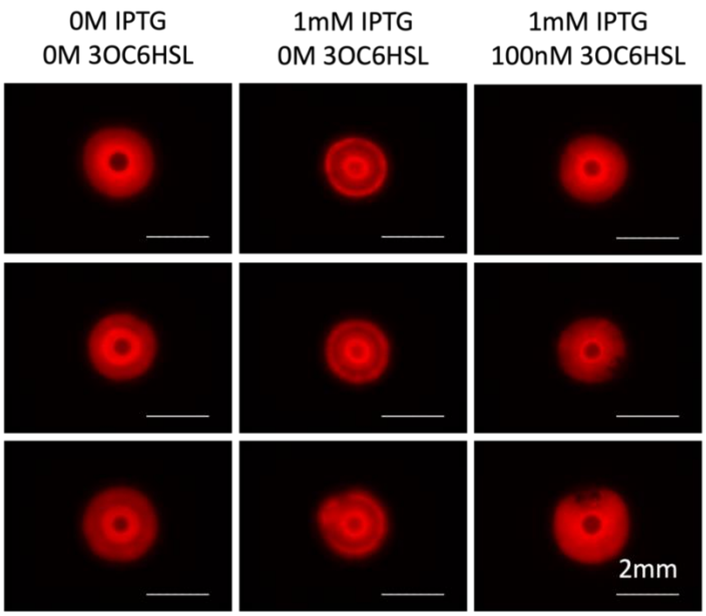


**Supplementary Figure 2.** IPTG and 3OC6HSL induction tunes pattern formation experimentally. 0.1 μl cell culture (OD = 0.2) was manually pipetted onto an agar petri dish containing the default growth medium. Combinations of 1 mM IPTG and 100 nM 3OC6HSL were added to the medium. Images were taken after 21 hours of growth. Adding IPTG enhanced ring formation; adding 3OC6HSL weakened ring formation.

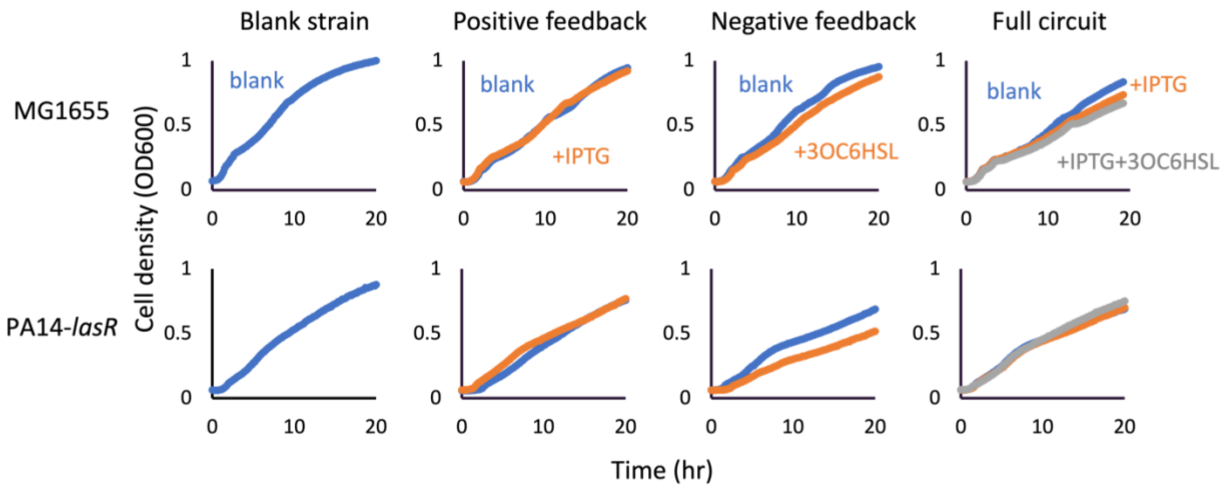


**Supplementary figure 3.** Growth curves of MG1655 and PA14-*lasR* carrying the patterning circuit in liquid growth medium. The growth medium contained 10g/L casamino acid and 10μg/ml gentamycin. 1mM IPTG, 100nM 3OC6HSL were added alone or together. The strains were either the blank host strain, the strain carrying the positive or negative feedback loop of the circuit, or the entire circuit (see plasmid list). MG1655 and PA14-*lasR* behaved alike in liquid culture.

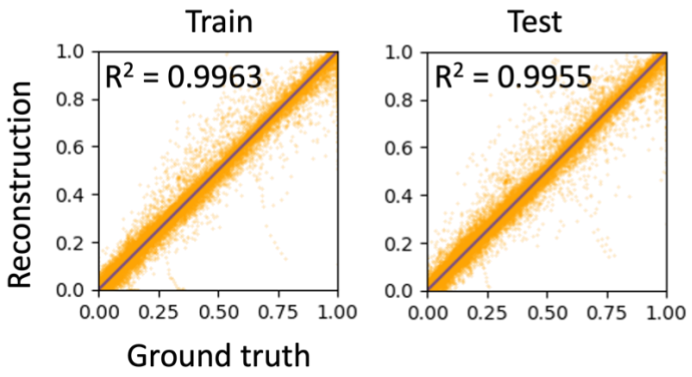


**Supplementary figure 4**. Variational autoencoder (VAE) training and test performance. The VAE was trained to reconstruct the 1D simulated mCherry profiles. It first compresses the signal into a lower dimensional space, then decompresses it back to the original dimensionality to reconstruct the original profiles. To demonstrate the model performance, 2000 random samples were selected from training and test sets, and $R^2$ between the ground-truth values (of 1D spatial profiles) and reconstructed values was used to evaluate the reconstruction accuracy. The blue line represents perfect alignment, $y = x$.

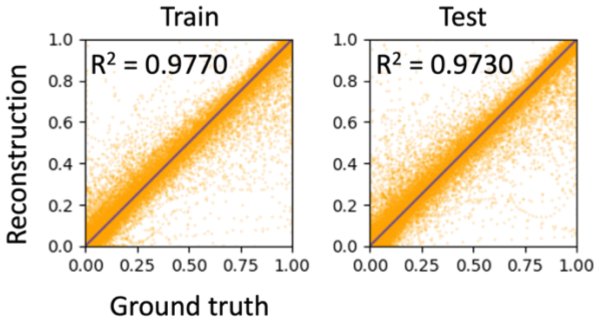


**Supplementary figure 5.** Multilayer perceptron – variational autoencoder (MLP-VAE) training and test performance. In the second step of the ML surrogate training, the VAE decoder was frozen, and an MLP was placed in front of it. The MLP was trained to predict the VAE latent variables using the 12 PDE parameters as inputs. These predicted latent variables were then passed through the VAE decoder to generate the 1D mCherry profile. A reconstruction loss of the 1D spatial profile was used to update MLP weights during training. To demonstrate the model performance, 2000 random samples were selected from training and test sets, and $R^2$ was used to evaluate the reconstruction accuracy. The blue line represents perfect alignment, $y = x$.

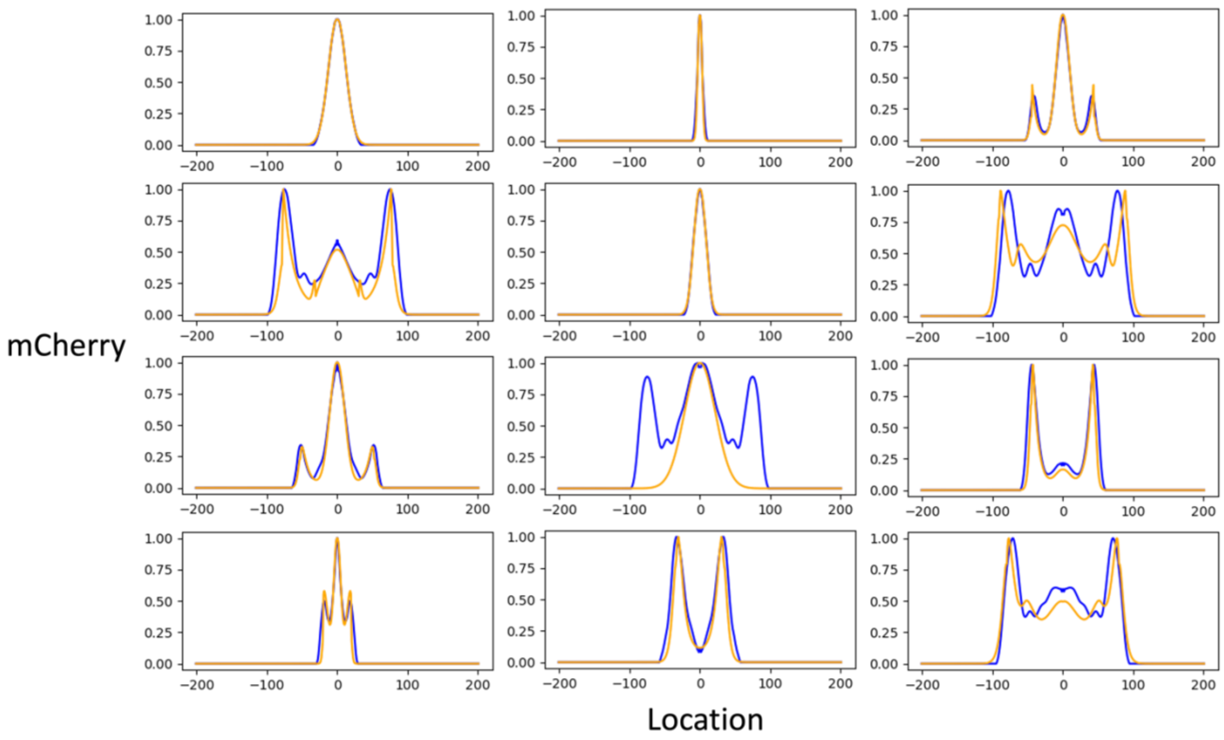


**Supplementary figure 6.** Validating ML predictions against PDE simulation. Most of ML predicted parameter sets match the PDE solutions, in terms of overall shape and formation of peaks, despite small quantitative differences. A small fraction shows significant deviation between ML and PDE simulations. Blue: ML inference, orange: PDE simulation.

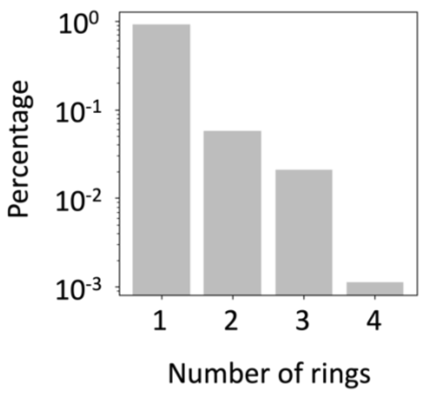


**Supplementary figure 7.** Pattern type distribution from ML-aided parameter screening. The parameter space exponentially shrinks with the number of rings. The total number of parameter combinations explored by the trained MLP-VAE model was 16million.

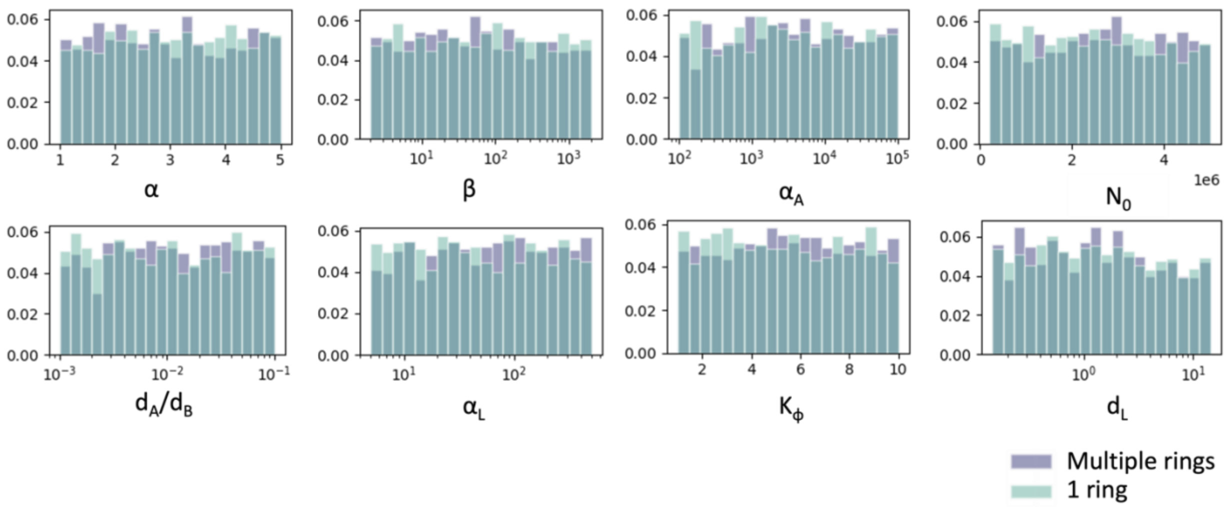


**Supplementary figure 8.** Parameters that were not critical for patterning. The analysis used the 4000 validated ML samples. These parameters have p-values from the Mann-Whitney U test greater than 0.05, indicating that circuit burden ($\alpha$, $\beta$), AHL synthesis rates ( $\alpha_A$), AHL degradation rates ($d_A$, $d_B$), lysozyme synthesis rate ($\alpha_L$), lysozyme degradation rate ($d_L$), gene expression profile ($K_\varphi$), and initial nutrient concentration ($N_0$) are not critical for multiple ring formation. For each of these parameters, the distributions of their values from the two groups (1 ring or multiple rings) are not statistically different.

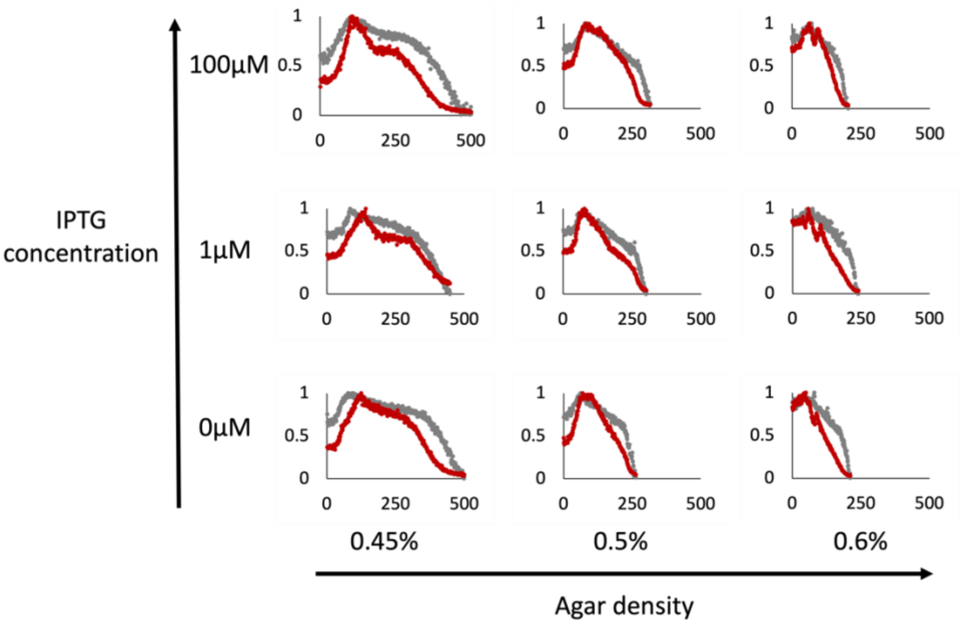


**Supplementary figure 9.** Experimental pattern formation with varied IPTG concentration and agar density (w/v). 0.1 µl cell culture (OD = 0.2) was printed ono the center of an agar dish containing 10 g/L casamino acid. The plates were incubated for 24 hours before imaging. Agreeing with the correlation found from computation, increasing IPTG and decreasing agar density synergistically enhanced multiple ring formation. For high agar density, increasing IPTG did not promote ring formation. Grey: brightfield, red: mCherry, both profiles are normalized with respect to their maximum values.

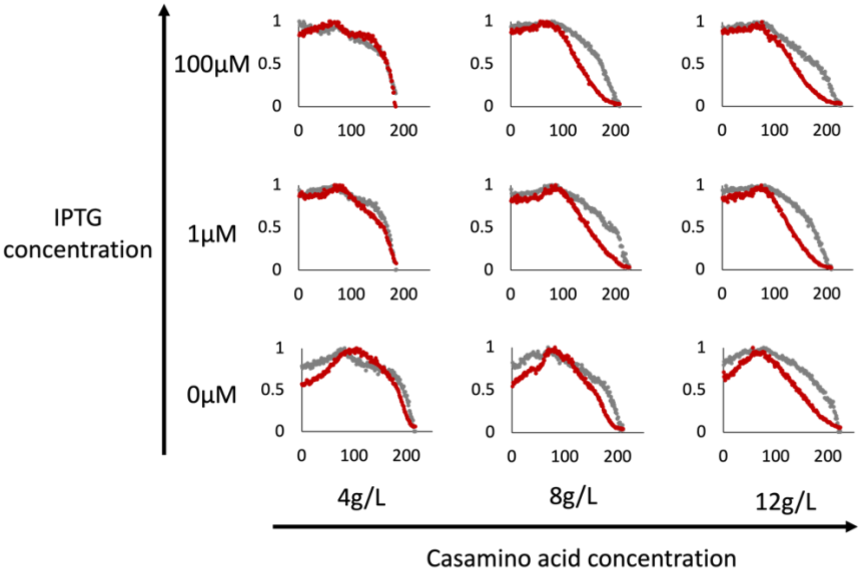


**Supplementary figure 10.** Experimental pattern formation with varied IPTG and casamino acid concentrations. 0.1 µl cell culture (OD = 0.2) was printed ono the center of an agar dish containing 0.5% agar (w/v). The plates were incubated for 24 hours before imaging. Agreeing with the correlation found from computation, increasing IPTG and decreasing growth rate (through lowering casamino acid concentration) synergistically enhanced multiple ring formation. For larger growth rates, adding IPTG did not lead to more rings. Grey: brightfield, red: mCherry, both profiles are normalized with respect to their maximum values.

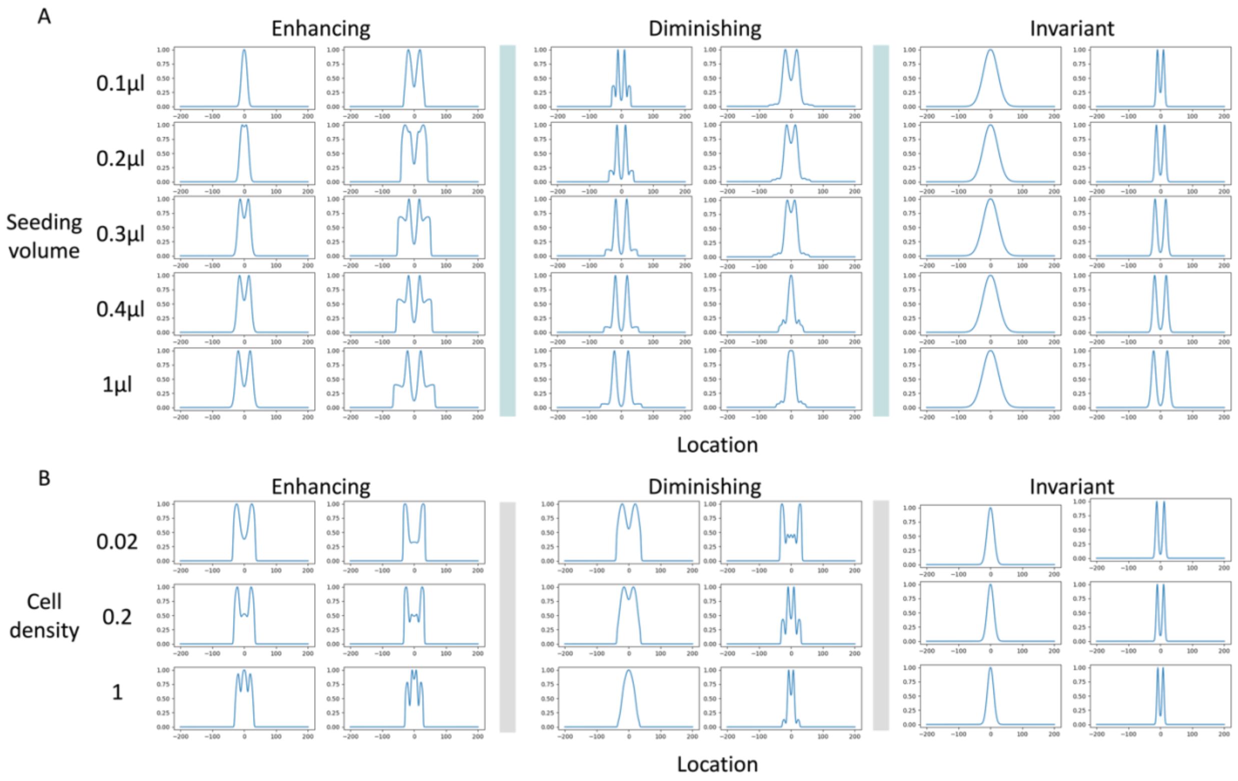


**Supplementary figure 11.** Effects of seeding volume and seeding cell density on pattern formation.

A. Computational screening on seeding volume. 3,000 random parameter sets were screened, where the original 12 PDE parameters were randomly sampled as before. For each parameter combination, the cell density was fixed at 0.2, and 5 seeding volumes (0.1, 0.2, 0.3, 0.4, 1 µl) were tested. Three distinct trends are identified: patterns either become more pronounced (e.g., more rings or deeper grooves), less pronounced (e.g., fewer rings or shallower grooves), or remain invariant to changes in seeding volume. Two examples from each category are presented.

B. Computational screening on cell density. 3,000 random parameter sets were screened, where the original 12 PDE parameters were randomly sampled as before. For each parameter combination, the seeding volume was fixed at 0.1 µl, and 3 cell densities (0.02, 0.2, 1) were tested. Three distinct trends are identified: patterns either become more pronounced (e.g., more rings or deeper grooves), less pronounced (e.g., fewer rings or shallower grooves), or remain invariant to changes in cell density. Two examples from each category are presented.

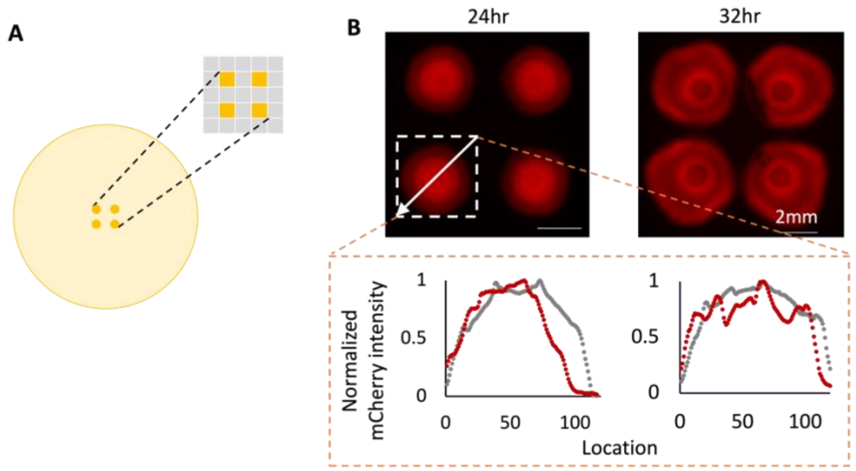


**Supplementary figure 12.** Pattern formation in colony arrays.

A. Seeding array. A 0.1 µL aliquot of cell culture (OD = 0.2) was printed into a 2×2 array at the center of a one-well plate agar dish. Each inoculum has a diameter of 0.88 mm, and the spacing between the centers of adjacent seeding positions is 4.5 mm (twice as far as in Figure 5D).

B. Experimental patterns from the colony array. Cross profiles were measured diagonally across the colonies, from the top right to the bottom left corner, passing through the colony centers along the white arrow. At 24 hours, the colonies formed a mCherry core. By 32 hours, asymmetric rings formed within each colony, with the rings closer towards the center. When placed apart, colonies did not branch out or merge. Grey: brightfield, red: mCherry, both are normalized with respect to their maximum values.
